# Macrophage proliferation machinery leads to PDAC progression, but susceptibility to innate immunotherapy

**DOI:** 10.1101/2021.11.08.467770

**Authors:** Chong Zuo, John M. Baer, Brett L. Knolhoff, Jad I. Belle, Xiuting Liu, Graham D. Hogg, Christina Fu, Natalie L. Kingston, Marcus A. Brenden, Angela Alarcon De La Lastra, Paarth B. Dodhiawala, Cui Zhou, C. Alston James, Li Ding, Kian-Huat Lim, Ryan C. Fields, William G. Hawkins, Guoyan Zhao, Jason D. Weber, David G. DeNardo

**Affiliations:** Department of Medicine, Washington University School of Medicine, St. Louis, MO 63110, USA; Siteman Cancer Center, Washington University School of Medicine, St. Louis, MO 63110, USA; Department of Surgery, Washington University School of Medicine, St. Louis, MO 63110, USA; Department of Pathology and Immunology, Washington University School of Medicine, St. Louis, MO 63110, USA; Department of Biology, Grinnell College, Grinnell, IA 50112, USA; Department of Cell Biology & Physiology, Washington University School of Medicine, St. Louis, MO 63110, USA; Department of Neuroscience, Washington University School of Medicine, St. Louis, MO 63110, USA

## Abstract

Tumor-associated macrophages (TAMs) are involved in many aspects of cancer progression and correlate with poor clinical outcomes in many cancer types, including pancreatic ductal adenocarcinomas (PDACs). Previous studies have shown that TAMs can populate PDAC tumors not only by monocyte recruitment but also by local proliferation. However, the impact local proliferation might have on macrophage phenotype and cancer progression is unknown. Here, we utilized genetically engineered cancer models, single-cell RNA-sequencing data, and *in vitro* systems to show that proliferation of TAMs was driven by colony stimulating factor-1 (CSF1) produced by cancer-associated fibroblasts. CSF1 induced high levels of p21 in macrophages, which regulated both TAM proliferation and phenotype. TAMs in human and mouse PDACs with high levels of p21 had more inflammatory and immunosuppressive phenotypes. The p21 expression in TAMs was induced by both stromal interaction and/or chemotherapy treatment. Finally, by modeling p21 expression levels in TAMs, we found that p21-driven macrophage immunosuppression *in vivo* drove tumor progression. Serendipitously, the same p21-driven pathways that drive tumor progression, also drive response to CD40 agonist. These data suggest that stromal or therapy-induced regulation of cell cycle machinery can regulate both macrophage-mediated immune suppression and susceptibility to innate immunotherapy.

**Summary:** TAMs are indicative of poor clinical outcomes and in PDAC their number is sustained in part by local proliferation. This study shows that stromal desmoplasia drives local proliferation of TAMs, and induces their immunosuppressive ability through altering cell cycle machinery, including p21 expression. Serendipitously, these changes in p21 in TAMs also potentially render tumors more sensitive to CD40 agonist therapy.

## Introduction

Macrophages are one of the most abundant immune cell types in the tumor microenvironment (Noy and Pollard, 2014). Extensive studies have shown that macrophages can mediate tumor immunosuppression by both directly interacting with cytotoxic T cells and indirectly affecting T cell functions through secretions of immuno-modulators that create a favorable tumor microenvironment (DeNardo and Ruffell, 2019; Cassetta and Pollard, 2018; Doedens et al., 2010). Aside from their immunosuppressive phenotypes, macrophages are known to promote tumor initiation, angiogenesis, local invasion, and metastatic spread (Ruffell and Coussens, 2015; Hao et al., 2012; Cassetta and Pollard, 2018). Unsurprisingly, the presence of macrophages is found to be associated with a poor clinical outcome in many cancers, including pancreatic cancer (Cassetta and Pollard, 2018; Ino et al., 2013). As such, preclinical and clinical studies have focused on targeting tumor-associated macrophages (TAMs). These approaches, often consisting of macrophage-depleting strategies, have yet to show clinical success, in spite of showing efficacies in preclinical models (DeNardo and Ruffell, 2019; Cannarile et al., 2017; Poh and Ernst, 2018; Xiang et al., 2021). This suggests more studies are needed to understand the varied subset of macrophages in tumors and how they impact tumor immunity and cancer progression.

During tissue damage, macrophage numbers can be increased by multiple mechanisms. These include the expansion of tissue resident macrophage populations by local proliferation or new macrophages can be recruited from blood monocytes (Ginhoux and Guilliams, 2016). This balance is likely regulated by both the tissues and types of damage. In pancreatic ductal adenocarcinoma (PDAC), macrophages are derived from both monocyte and tissue resident sources (Zhu et al., 2017). One consistent characteristic of TAMs from both sources in PDAC mouse models is that they are highly proliferative (Zhu et al., 2017). Notably, proliferation of macrophages is not only observed in tumors, but also in injured and inflamed tissues (Hashimoto et al., 2013; Davies et al., 2011; Van Gassen et al., 2015). Under these conditions, inhibiting macrophage proliferation dramatically reduced macrophage number and inflammation (Tang et al., 2015). These observations raised the possibility that inhibiting macrophage proliferation in PDAC might limit the number of tumor-promoting macrophages.

Macrophage proliferative status is commonly associated with underlying macrophage phenotypes. Interferon gamma (IFN-*γ*) and lipopolysaccharide (LPS) inhibit macrophage proliferation and induce production of nitric oxide (NO) and inflammatory cytokines (Müller et al., 2017; Xaus et al., 2000; Marchant et al., 1994). Interleukin (IL)-4 promotes macrophage proliferation and drives them to a T_H_-2 like phenotype (Jenkins et al., 2013). These observations led to the question of whether the macrophage proliferation machinery plays a role in regulating macrophage phenotypes.

In this study, we aimed to understand how the PDAC microenvironment drove local macrophage proliferation and what the net outcome of this was on tumor immunity and progression. We discovered that while cancer-associated fibroblast-induced macrophage proliferation was important for sustaining TAM number, induction of p21 in TAMs by stromal colony stimulating factor-1 (CSF1) resulted in immunosuppression and tumor progression.

## Results

### Tumor infiltrating macrophages are highly proliferative in PDAC

To evaluate human PDAC infiltration by TAMs, we utilized multiplex immunohistochemistry (mpIHC) to stain for CD68^+^ macrophages and CK19^+^ tumor cells in human PDAC tissues and found that CD68^+^ TAMs were more frequent in PDAC tissues when compared to adjacent normal pancreas tissues (Fig. 1 A). To further study infiltrating macrophages, we utilized a p48^−^Cre^+^/LSL-Kras^G12D^/p53^flox/flox^ (KPC) genetically engineered mouse model (GEMM), which spontaneously develops PDAC tumors and recapitulates the pathological features of human PDAC (Hingorani et al., 2003, 2005). As in human PDAC, we found that the number of F4/80^+^ TAMs increased paralleling disease progression (Fig. 1 B and Fig. S1 A). Our previous studies have shown that these PDAC infiltrating TAMs were sustained by both local proliferation and monocyte recruitment in animal models (Zhu et al., 2017). However, these studies did not assess the potential impact macrophage proliferation might have on tumor progression or tumor immunity.

**Figure 1.**
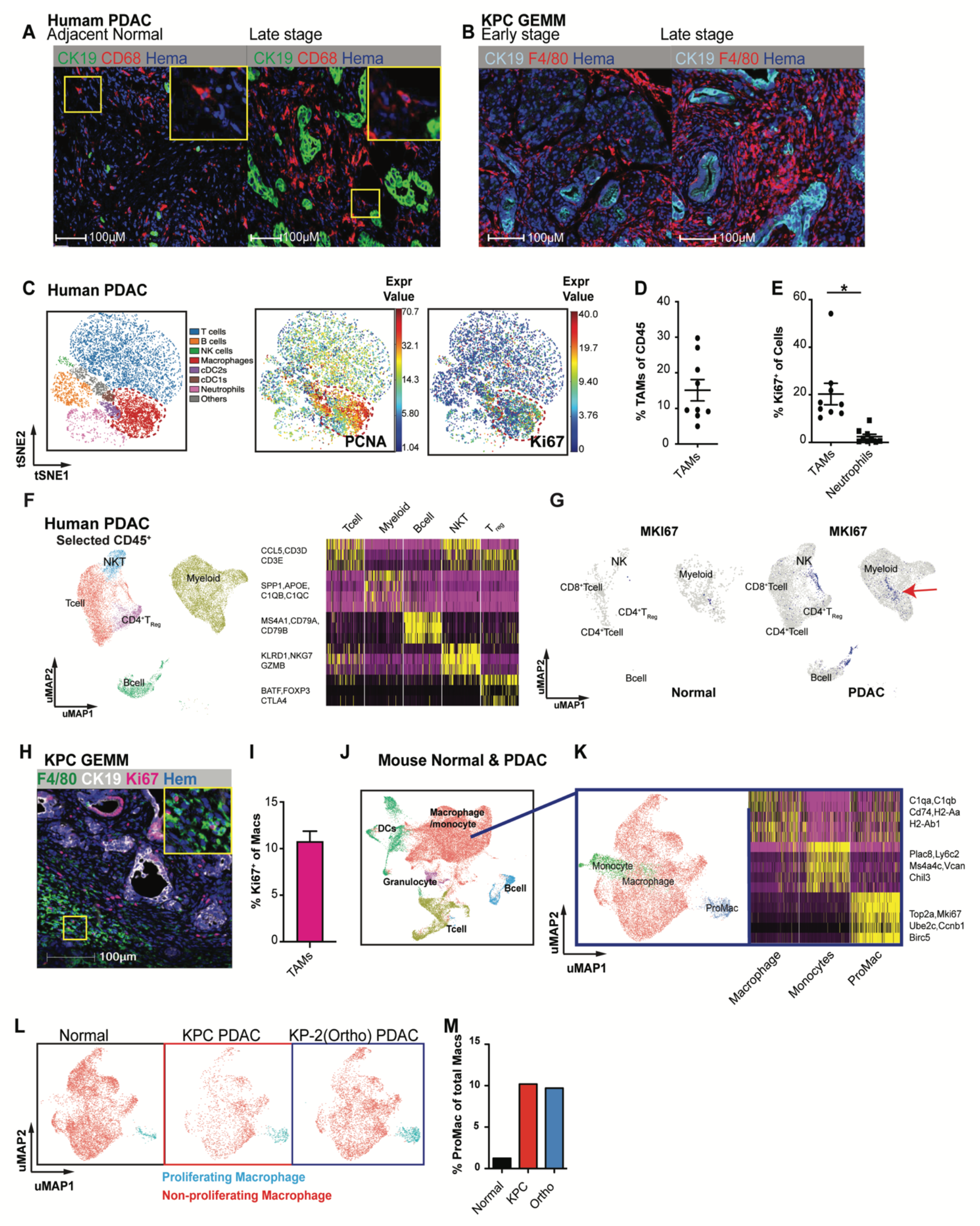
Pancreatic ductal adenocarcinoma (PDAC)-infiltrating macrophages are highly proliferative. **(A)** Representative immunohistochemistry (IHC) analyses of CD68^+^ macrophages and CK19^+^ tumor cells in the late stage of PDAC tissues and adjacent normal tissues from human patients. (**B)** Representative IHC analyses of F4/80^+^ macrophage and CK19^+^ tumor cells in early and late stages of KPC genetically engineered mouse models (GEMMs). **(C)** Representative tSNE plots of total normalized CD45^+^ cells from a PDAC patient, annotated with manually assigned cell identity. The macrophage cluster was marked with a red circle, and expressions of PCNA and Ki67 were explicitly displayed. **(D,E)** Dot plot displaying quantification of tumor-associated macrophages (TAMs), Ki67^+^ TAMs, and Ki67^+^ neutrophils across nine human PDAC patients. **(F)** UMAP of realigned and reprocessed publicly available human pancreatic ductal adenocarcinoma (PDAC) dataset (Peng et al., 2019) displaying major CD45^+^ clusters with expression levels of MKi67 and a heat map showing key gene expressions for each cluster. n = 21 PDAC samples, n = 6 normal samples. **(G)** UMAP plots displaying normalized expression levels of MKI67 across subpopulations with red arrow pointing to MKI67 expressing myeloid cells. **(H,I)** Representative multiplex immunohistochemistry (mpIHC) displaying F4/80^+^ macrophages, CK19^+^ tumor cells, and Ki67^+^ proliferating cells in tumors from p48^−^Cre^+^/LSL-Kras^G12D^/p53^flox/flox^ (KPC) GEMMs with quantification of Ki67^+^ macrophages; n = 6 mice. **(J)** UMAP dimensionality reduction plot of integrated sorted CD45^+^ cells from the murine normal pancreas and pancreatic tissues from KPC PDACs and KP-2 orthotopic PDACs with cell type annotations and cell cycle regression. **(K)** UMAP plot of reclustered macrophages/monocytes in **J** without cell cycle regression with a heat map displaying corresponding gene signatures. **(L)** UMAP displaying proliferating macrophages and non-proliferating macrophage clusters across the mouse scRNAseq data set used in **J** with quantification in **(M)**. Data are presented as the mean ± SEM. n.s., not significant; *p < 0.05. For comparisons between any two groups, Student’s two-tailed *t*-test was used.

To further investigate the significance and mechanisms of local proliferation of TAMs, we more deeply studied pancreatic tissues from GEMMs and human PDAC patients. We first evaluated the frequency of proliferating macrophages in human PDAC tumors by mass cytometry time of flight (CyTOF). Distinguishing major leukocyte populations based on surface markers, we found that CD68^+^CD64^+^ macrophages composed >15% of all infiltrating leukocytes (Fig. 1, C and D and Fig. S1 B). Notably, these macrophages expressed high levels of the proliferation markers PCNA and Ki67 (Fig. 1 C). Ki67^+^ macrophages made-up 20% of total macrophages, and this percentage was significantly higher than that of other leukocyte populations, such as neutrophils (Fig. 1 E and Fig. S1 C). Next, we examined proliferating macrophages in tumors from KPC GEMMs. We observed >10% of F4/80^+^ cells were also Ki67^high^ by mpIHC analysis (Fig. 1, H and I). In addition, we generated and analyzed single-cell RNA-sequencing (scRNAseq) data from normal pancreas, pancreatic tissues from KPC GEMMs, orthotopic PDAC tumors, and previously published human PDAC datasets (Peng et al., 2019) (Fig. S1 D). In human PDACs, we found populations carrying both myeloid and proliferating signatures (Fig. 1, F and G). Similarly, in mouse datasets, we identified TAMs independent of cell cycle genes (Fig. 1 J and Fig. S1 E), then upon reclustering, we easily identified discrete clusters with cell cycle gene signatures (Fig. 1 K). As expected, this cluster was expanded in PDACs compared to normal tissues (Fig. 1, L and M). Taken together, these data suggest that a significant portion of macrophages are actively proliferating in both murine and human PDAC tissues.

### Cancer-associated fibroblasts drive macrophage proliferation through CSF1

To identify the cellular players that drove macrophage proliferation in PDAC, we investigated the cellular composition in the PDAC tumor microenvironment (TME). As others have shown, PDAC tumors contain dense fibrotic stroma (Elyada et al., 2019; Schnittert et al., 2019; Waghray et al., 2013), and immunohistochemistry (IHC) staining of PDAC tissues from KPC GEMMs revealed abundant PDPN^+^ cancer-associated fibroblasts (CAFs) surrounding CK19^+^ tumor cells (Fig. 2 A). We next performed proximity analysis and found that TAMs were within 100 µm to both tumor cells and CAFs, but more frequently closer to PDPN^+^ CAFs than CK19^+^ tumor cells (Fig. 2 B). To test whether fibroblasts and tumor cells drove macrophage proliferation, we co-cultured bone marrow-derived macrophages (BMDMs) with either PDAC cell lines from KPC GEMMs or primary pancreatic fibroblasts. We found that PDAC cells and fibroblasts both led to increases in macrophage proliferation, as measured by BrdU incorporation. However, fibroblasts induced significantly higher levels of proliferation and increases in the number of macrophages (Fig. 2 C). Additionally, macrophage proliferation was not further enhanced by triple culture of PDAC cells and fibroblasts, suggesting the effects were not additive (Fig. 2 C, grey bars). To determine if fibroblasts induced macrophage proliferation in a cell contact-dependent manner or through secreted factors, we repeated these assays in a Transwell system. We found that without direct contact to BMDMs, fibroblasts still drove macrophage proliferation at almost a comparable level as the strong mitogen, CSF1 (Fig. 2 D).

**Figure 2.**
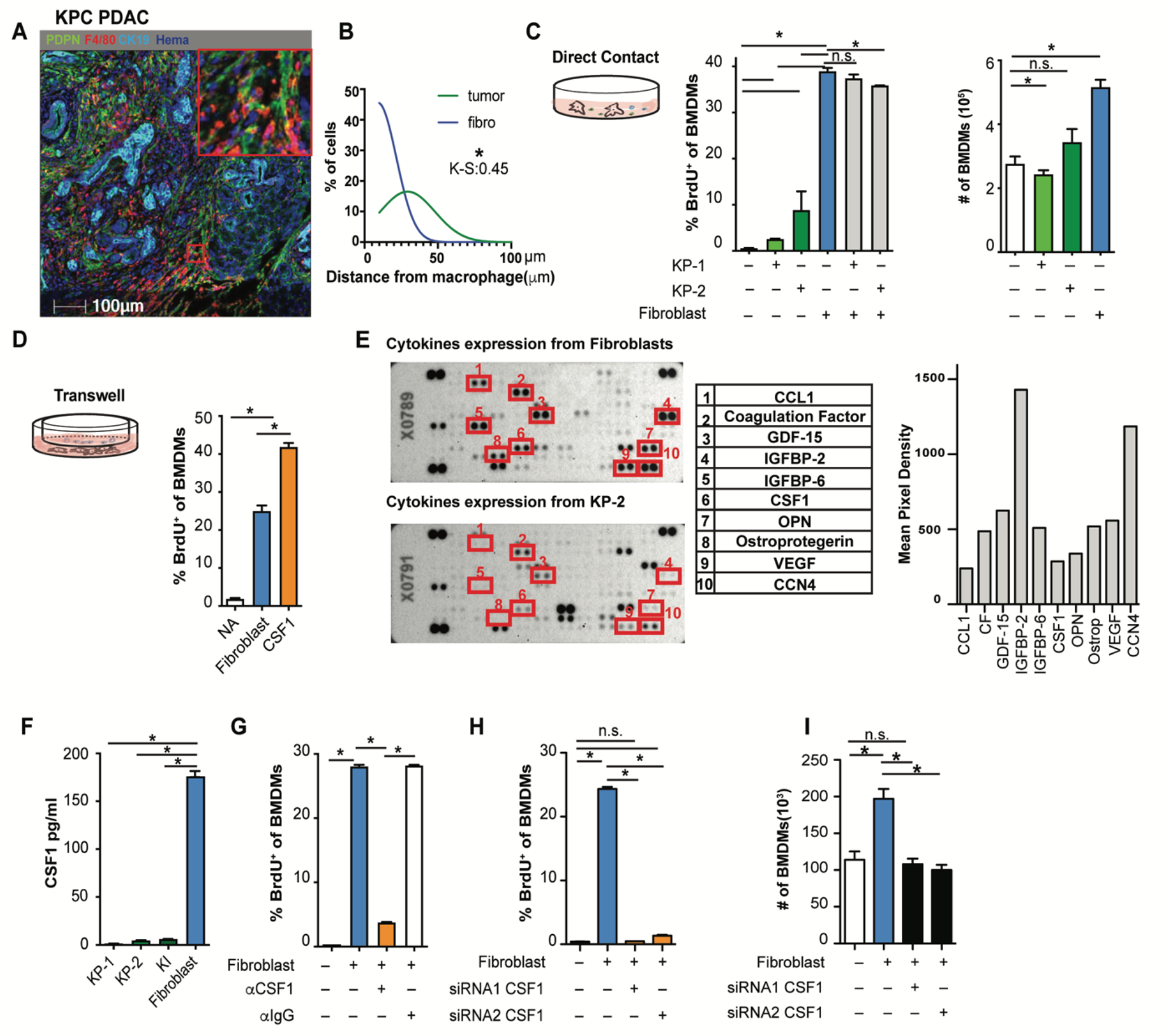
Fibroblasts drive macrophage proliferation through colony stimulating factor-1 (CSF1). **(A)** Representative multiplex immunohistochemistry (mpIHC) image of p48^−^Cre^+^/LSL-Kras^G12D^/p53^flox/flox^ (KPC) mouse pancreatic ductal adenocarcinomas (PDACs) displaying alpha smooth muscle actin (αSMA^+^) (white) fibroblasts, CK19^+^ (teal) tumor cells, and F4/80^+^ (green) macrophages. **(B)** Frequency distribution of Pdpn^+^ fibroblasts (blue curve) and CK19+ tumor cells (green curve) to a nearest F4/80^+^ macrophage. n = 6 KPC mice. **(C)** The 5-bromo-2’-deoxyuridine (BrdU) incorporation and number of bone marrow-derived macrophages (BMDMs) in co-culture with KP-1, KP-2, fibroblasts, or the combination for 48 h, BrdU pulsed for the last 6 h; n = 6. **(D)** The BrdU incorporation of BMDMs when cultured with fibroblasts in a Transwell assay or 10 ng/mL of CSF1 for 48 h, and BrdU pulsed for the last 6 h; n=3. **(E)** Representative image of a cytokine antibody array resulting from fibroblast- and KP-2-conditioned media, highlighting the top 10 highly expressed cytokines in fibroblast-conditioned medium and the corresponding mean pixel densities. The arrays were repeated two times. **(F)** Bar graph shows the concentrations of CSF1 from three tumor-conditioned media (KP-1, KP-2, and KI) and fibroblast-conditioned medium measured by an ELISA. **(G)** BrdU incorporation of BMDMs in co-culture with fibroblasts treated with 2 µg of *α*CSF1 or 2 µg of *α*IgG for 24 h, and BrdU pulsed for the last 6 h; n=3. **(H,I)** BrdU incorporation and number of BMDMs in Transwell cultures with fibroblasts with or without siRNA knockdown for CSF1; n=3. Data are presented as the mean ± SEM. n.s., not significant; *p<0.05. All *in vitro* assays were consistent across at least two independent repeats. For comparisons between any two groups, Student’s two-tailed *t*-test was used. Frequency distributions were compared using the nonparametric Kolmogorov-Smirnov test.

To identify the relevant secreted factors from fibroblasts that drove macrophage proliferation, we profiled 111 soluble factors derived from two PDAC cell lines (KP-1, KP-2), or fibroblast-conditioned media and found that fibroblasts secreted significantly higher levels of CSF1 (Fig. 2 E). We measured the levels of CSF1 secreted by fibroblasts and three different PDAC cell lines (KP-1, KP-2, and KI) through ELISAs and confirmed that only fibroblasts produced high levels of CSF1 (Fig. 2 F). Next we sought to determine if CSF1 was necessary and sufficient for fibroblasts to drive macrophage proliferation. Both the addition of neutralizing *α*CSF1 IgG to the co-culture of BMDMs and fibroblasts, and knocking-down CSF1 in fibroblasts by siRNA in Transwell assays, resulted in a loss of fibroblast-driven macrophage proliferation and number expansion (Fig. 2, G,H and I). These data suggest that CSF1 secreted from fibroblasts is both necessary and sufficient for macrophage proliferation *in vitro*.

To confirm CAFs drive TAMs proliferation in *in vivo* pancreatic tissue, we analyzed scRNAseq datasets from both mouse and human. In a previously published dataset (Hosein et al., 2019) of pancreatic tumors from three GEMM models, including *Kras^LSL-G12D/+^Ink4a^fl/fl/^Ptf1a^Cre/+^ (KIC), Kras^LSL-G12D/+^Trp53^LSL-R172H/+^Ptf1a^Cre/+^ (KP^R172H/+^C),* and *Kras^LSL-G12D/+^Trp53^fl/fl/^Pdx1^Cre/+^* (*KPfC*), we found that fibroblasts expressed higher levels of CSF1 than other cell types (Fig. 3, A and B). In a human PDAC dataset (Peng et al., 2019) comprised of 21 PDAC samples, fibroblasts also expressed a higher level of CSF1 than tumor cells and other cells within the TME (Fig. 3, C and D). Others have also detected CSF1 in the cultures of primary CAFs from PDAC patients (Samain et al., 2021). Collectively, these data suggest that fibroblasts are the main producers of CSF1 in the PDAC TME. Next, we injected *α*CSF1 IgG into mice bearing orthotopic KP-2 tumors and measured macrophage proliferations 12 and 24 h after the injection. Similar to the *in vitro* experiments, we found a significant reduction in the percentage of macrophages undergoing proliferation, measured by BrdU incorporation (Fig. 3, E, F and G). We have previously shown that sustained CSF1 depletion, exceeding 48 h, led to macrophage depletion by apoptosis (Zhu et al., 2014). To eliminate the possibility that the decrease in proliferation came from macrophage death, we quantified macrophage numbers and found no change (Fig. 3 H). Additionally, we found that proliferation of monocytes was minimal and not significantly affected by *α*CSF1 IgG treatment (Fig. 3 I), confirming that the reduction of proliferation was mainly from macrophages. Taken together, these data suggest that CSF1 secreted by cancer-associated fibroblasts drives local macrophage proliferation in pancreatic cancer.

**Figure 3.**
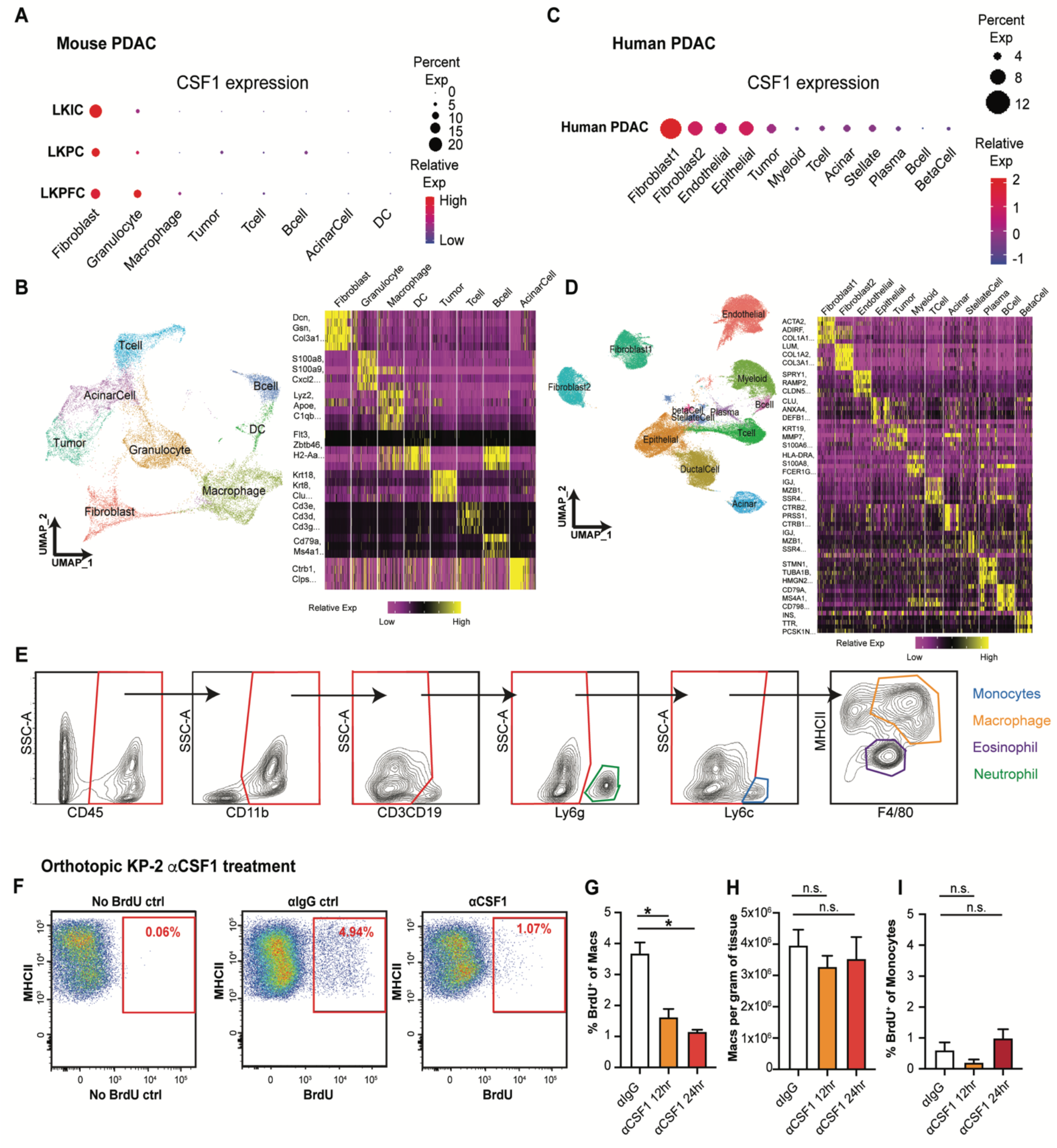
Cancer-associated fibroblasts drive tumor associated macrophage proliferation through colony stimulating factor-1 (CSF1). **(A)** Dot plot summarizing CSF1 expressions in different cell types across three mouse PDAC models from the publicly available scRNAseq dataset (Hosein et al., 2019). **(B)** UMAP dimensionality reduction plot of integrated cells from LKIC, LKP *^R172H/+^*C, and LKPFC genetically engineered mouse models in scRNAseq dataset used in **A**, annotated with different cell types. Data were filtered and reprocessed as described in the **Methods**. **(C)** Dot plot displaying CSF1 expressions in different cell types across 21 human PDAC patient samples from the publicly available scRNAseq dataset (Peng et al., 2019). **(D)** UMAP dimensionality reduction plot of integrated cells from 21 pancreatic adenocarcinoma patients used in **C**, annotated with different cell types. **(E)** Representative flow cytometry plots showing the gating strategy to identify macrophages, monocytes, neutrophils in orthotopic KP-2 tumors. **(F-I)** Representative flow cytometry plot and quantification bar plot showing BrdU^+^ macrophages and monocytes, and total number of macrophages following *α*IgG or *α*CSF1 injections; n = 6-8 mice per group. Data are presented as the mean ± SEM. n.s., not significant; *p<0.05. For comparisons between any two groups, Student’s two-tailed *t*-test was used.

### The p21 cell cycle-dependent kinase inhibitor was induced in TAMs by CAF-derived CSF1

We next asked whether the macrophage proliferation machinery regulated by CAF-derived CSF1 could impact the TAM phenotype. We first examined the expressions of several critical cell cycle regulators in BMDMs following treatment with either CSF1, the proliferative mitogen, or lipopolysaccharide (LPS), which is known to blunt macrophage proliferation (Liu et al., 2016) (Fig. S2 A). We found that when BMDMs were treated with CSF1, overall protein levels of c-Myc and cyclin D1 were upregulated while p27^Kip1^ was reduced (Fig. 4 A). BMDMs treated with LPS showed the opposite result. These changes are consistent with the existing roles of cell cycle promoters (c-Myc and cyclin D1) and a cell cycle inhibitor (p27^Kip1^) (Liu et al., 2016; Matsushime et al., 1991). However, surprisingly, we found p21^Waf/Cip1^, a cell cycle inhibitor (Cazzalini et al., 2010; Dutto et al., 2015; Brugarolas et al., 1999), was strongly induced by both CSF1 and CAF co-culturing (Fig. 4, B and D). To further investigate this p21 induction, we performed a kinetic study of p21 expression in BMDMs and found that the p21 protein was induced by CSF1 within 6–12 h, which was prior to S phase entry at 24–48 h after CSF1 administration, as measured by BrdU (Fig. 4, B and C). Similar kinetics and cell cycle transit were found when BMDMs were cultured with fibroblasts in a Transwell assay (Fig. 4, D and E). These data suggest that p21 induction by stoma-derived CSF1 could impact both macrophage cell cycle and phenotype.

**Figure 4.**
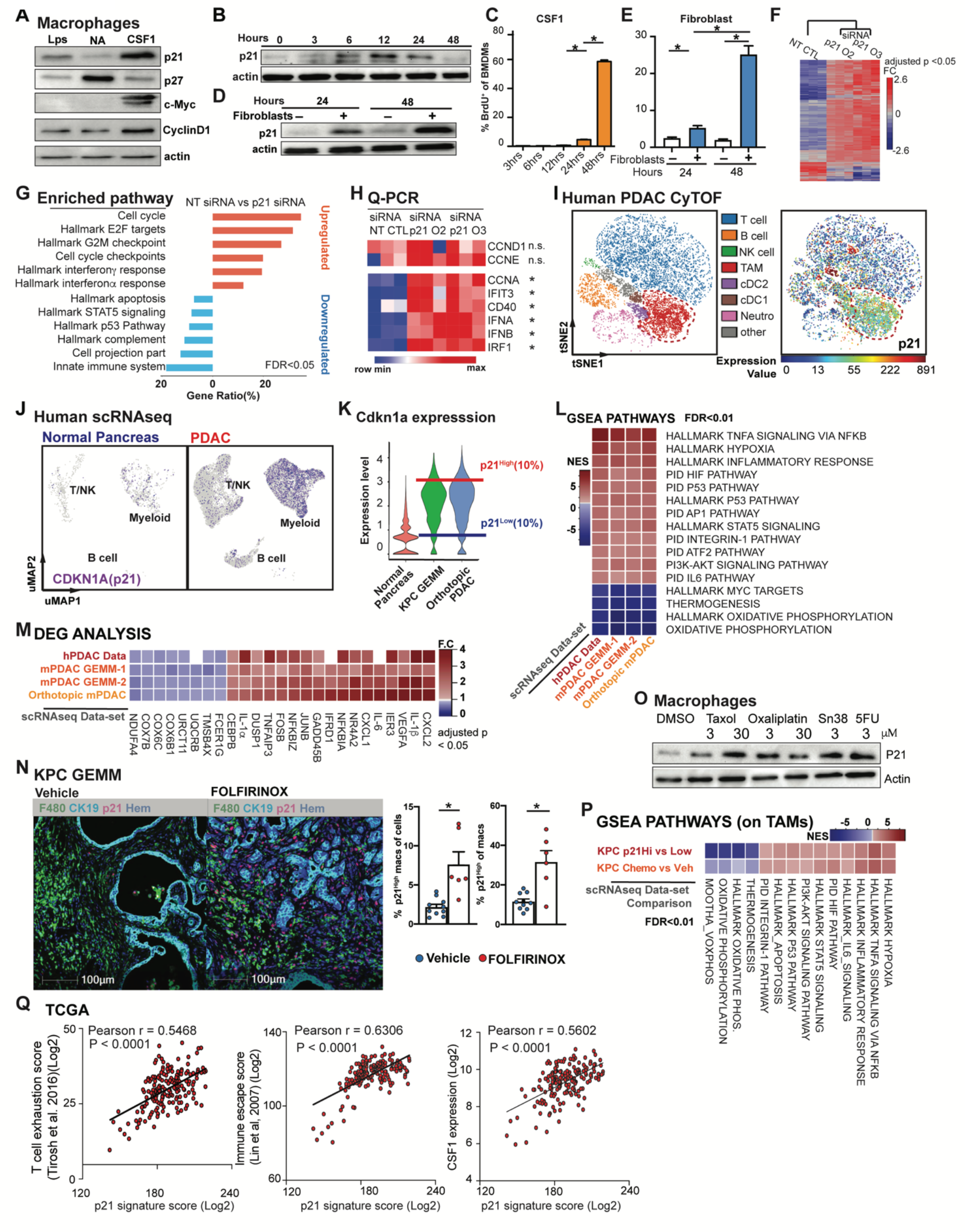
The p21 cell cycle-dependent kinase inhibitor is induced by CSF1 and regulates the macrophage phenotype. **(A)** Immunoblots of p21, p27, c-Myc, and cyclinD1 in bone marrow-derived macrophages (BMDMs) after treatment with 100 ng/mL of lipopolysaccharide or colony stimulating factor-1 (CSF1) for 24 h. The experiments were repeated three times. **(B)** Immunoblot displaying p21 expression in BMDMs following 4 ng/mL CSF1 treatment at time 0 with quantification of BrdU^+^ BMDMs shown in **(C)**, 5-bromo-2’-deoxyuridine (BrdU) was added at time 0 and pulsed until harvest. BMDMs were starved without CSF1 overnight. **(D)** Immunoblot displaying p21 expression in BMDMs combined with fibroblasts in Transwell assays at time 0. **(E)** Bar plot displaying the quantification of BrdU^+^ BMDMs in **D**. **(F)** Heat map displaying the microarray analysis of differentially expressed genes (DEGs) between non-target siRNA treated or siRNA targeting for p21 treated BMDMs cultured in tumor-conditioned medium for 24 h; n = 3 per group. Genes were filtered with adjusted p < 0.05 and fold-change > or < 1.5. **(G)** Bar graph displaying top overrepresentation analysis of DEGs in **F** to known biological functions [Gene Ontology (GO), Kyoto Encyclopedia of Genes and Genomes (KEGG), REACTOME, and Molecular Signatures Database (MSigDB)] with a false discovery rate (FDR) < 0.05. **(H)** Heat map displaying qPCR analysis of gene expressions of cell cycle and interferon-related genes between non-target siRNA treated or siRNA targeting for p21 treated BMDMs cultured in tumor-conditioned medium for 24 h; fold-change > 1.5, n = 3/group of the comparison. **(I)** Representative tSNE plot displaying major cell types from CyTOF analysis of a human PDAC patient (same as in **Fig. 1 C**) with macrophages circled in red and p21 expression. **(J)** UMAP displaying CDKN1A gene expression in CD45^+^ cells from the human PDAC scRNAseq dataset (Peng et al., 2019) with annotation of key cell types. **(K)** Violin plot showing the expression levels for p21 gene in macrophage clusters from integrated scRNAseq analyses of the mouse normal pancreas and pancreatic tissue from KPC GEMMs and orthotopic KP-2 tumor-bearing mice. Representative lines were drawn for two groups of stratified macrophages based on the top 10% of p21 expression and bottom 10% of p21 expression. **(L)** Heat map of net enrichment score (NES) of shared enriched pathways identified by GSEA analysis comparing the two groups of macrophages (p21^High^ vs. p21^Low^) in human PDAC scRNAseq dataset (23), (27), KPC GEMM and orthotopic scRNAseq data. Enriched pathways were selected by FDR < 0.01. **(M)** Heat map displaying the shared DEGs when comparing p21^High^ to p21^Low^ tumor-associated macrophages (TAMs) in each dataset with adjusted p < 0.05 and fold-change > 1.2 or < 0.8. p21^High^ signature score was created utilizing filtered DEGs with fold-change > 1.5 across three mouse scRNAseq datasets. **(N)** Representative mpIHC image displaying F4/80^+^ TAMs, CK19^+^ tumor cells, and p21^+^ cells in KPC GEMM treated with dimethyl sulfoxide or FOLFIRINOX for 24 h with quantification of p21^+^TAMs as total cells and total TAMs on the right. **(O)** Immunoblots showing expressions of p21 in BMDMs after treatment with chemotherapeutics for 24 h. **(P)** Heat map of NES of shared enriched pathways identified by GSEA analysis in comparing p21^High^ to p21^Low^ TAMs in KPC GEMM PDAC and in comparing chemotherapeutic treated KPC GEMM PDAC to DMSO treated KPC GEMM PDAC with FDR < 0.05. **(Q)** Correlation plots with Pearson coefficients (r) of p21 signature score vs. T cell exhaustion score (Tirosh et al., 2016), Immune escape score (Lin et al., 2007), and CSF1 expression from TCGA PDAC PanCancer Atlas study (n=180). All graphs are expressed as the mean ± SEM. n.s., not significant; *p < 0.05. All *in vitro* assays and immunoblots were consistent across more than two independent repeats. For comparisons between any two groups, Student’s two-tailed *t*-test was used, except for **F, M** where the Bonferroni correction was used and for **L, P** where the FDR was used.

To test if p21 induction impacted macrophage phenotype, we knocked-down p21 expression in BMDMs by siRNA in the presence of CSF1. We found that p21 knockdown resulted in a significant increase in the number of macrophages that entered S phase, confirming p21’s inhibitory role in the G1/S transition (Fig. S2 B and C). To assess macrophage phenotypic changes after p21 knockdown, we performed gene profiling analysis followed by RT-qPCR validation of altered gene expressions. Transcription profiling revealed > 300 genes that were differentially expressed in BMDMs upon p21 knockdown in the presence of tumor conditioned medium (Fig. 4 F). Overrepresentation analysis of the differentially expressed genes demonstrated that p21 knockdown in BMDMs resulted in the upregulation of genes involved in cell cycle progression, as expected, but also unexpectedly, it upregulated interferon *α* and *γ* responses (Fig. 4 G). RT-qPCR validation also found upregulation of interferon-related genes, IFIT3, CD40, IFN-*α* and IFN-*β*. Notably, gene expression of cyclins involved in early cell cycle stage (G1), CCND1, CCNE, were unchanged, while CCNA, an S phase cyclin, was upregulated (Fig. 4 H). Together, these data suggest that in addition to its canonical role in regulating S phase entry, p21 might suppress interferon signaling pathways. In a CSF1-rich TME like PDAC, elevated p21 expression in macrophages might play a prominent role in impairing tumor immunity (Hervas-Stubbs et al., 2011).

Based on the significant presence of CSF1-producing CAFs in the PDAC TME, we hypothesized that p21 might be chronically high in TAMs and thus might drive their immune-suppressive phenotype. We first evaluated p21 expression in human PDAC tumors by CyTOF, and found PDAC TAMs frequently expressed high levels of p21 (Fig. 4 I). Similarly, KPC tumors also had significant numbers of F4/80^+^ TAMs expressing high levels of p21 evaluated by mpIHC (Fig. S2, H and I). Finally, scRNAseq analysis suggested that TAMs from both human and murine PDAC tissues had higher levels of p21 gene expression than macrophages in normal tissues (Fig. 4 J; Fig. S2 F). The elevation of p21 in PDAC tumors could be a result of increased number of macrophages entering cell cycle as shown in Figure 1, G and L. However, we observed in CyTOF, TAMs that were high in p21 expression, were not necessarily high in the expression of PCNA or Ki67 (Fig. 4 I; Fig. 1 C), suggesting p21 expression was not only up in proliferating TAMs. In addition, we did not find a significant difference in the p21 protein levels between Ki67^+^ vs. Ki67^−^ TAMs by CyTOF, nor did we find significant difference in p21 gene expression in proliferating and non-proliferating clusters of TAMs in scRNAdeq data (Fig. S2, D and G). Collectively, these results suggest that elevated p21 expression in PDAC TAMs is unlikely to be solely caused by cell cycle entry/progression, it may become elevated by other factors in the TME and regulate TAMs phenotype.

To further assess the potential phenotypic differences in TAMs based on p21 expression, we generated and analyzed data from four scRNAseq data sets, including one from human (Peng et al., 2019) and three from PDAC mouse models (Hosein et al., 2019). We identified macrophage populations in each mouse dataset and myeloid populations in human dataset based on known macrophage markers after unsupervised clustering and UMAP projection (Fig. 1, F and J; Fig. 3 B). We then stratified macrophages (myeloid cells in human) based on p21 gene expressions to the p21^High^ and p21^Low^ grouped in each data set (Fig. 4 K). Notably, UMAP dimension reduction revealed the similar spatial distributions of p21^High^ and p21^Low^ macrophages in tumors from mouse GEMM and orthotopic models, suggesting shared characteristics among the same group of TAMs in different models (Fig. S2 E). To understand what these common phenotypes were, we performed Gene Set Enrichment Analysis between p21^High^ and p21^Low^ macrophages in each dataset. Across all four datasets and both species, we found that hallmarks typically associated with the tumor necrosis factor alpha (TNF-*α*) signaling pathway, hypoxia, and STAT5 signaling were upregulated in p21^High^ macrophages (p21^High^ myeloid cells in human), while oxidative phosphorylation pathways were upregulated in p21^Low^ macrophages (p21^Low^ myeloid cells in human) (Fig. 4 L). Although TNF-*α* and its signaling pathway are proinflammatory, they are frequently considered immunosuppressive in tumors. In this respect, TNF-*α* can mediate T cell exhaustion, CD8^+^ T cell death, and expansion of myeloid-derived suppressor cells and regulatory T cells (T_Regs_) to promote tumor progression and metastasis (Salomon et al., 2018; Balkwill, 2006). Consistent with the enrichment for TNF-*α* via the NF-*κ*B signaling pathway, expressions of IL-1*α*, IL-1*β*, and NF-*κ*B components were also upregulated in p21^High^ macrophages (Fig. 4 M). Together, these data suggest that TAMs with high p21 expression acquire an inflammatory but potentially immunosuppressive gene signature.

PDAC patients are frequently treated with cytotoxic chemotherapies, that can impact both tumor cells as well as stromal cells. Therefore, we sought to next determine if chemotherapy could impact TAM proliferation and p21 expression and thus influence TAM-immunosuppressive programs. First, we treated KPC GEMM with modified FOLFIRINOX (5-FU, Irinotecan, and Oxaliplatin), and analyzed p21^High^F4/80^+^ TAMs 24 hours later by mpIHC. We found that the number of p21^High^ TAMs significantly increased after chemotherapy treatment (Fig. 4 N). To determine if this was a direct effect of chemotherapeutic exposure, we treated BMDMs with four different chemotherapeutics for 24 h and observed similar inductions of p21 (Fig. 4 O). Finally, to assess if this induction of p21 by chemotherapy correlates with changes in macrophage phenotype, we analyzed TAMs from KPC GEMMs treated with vehicle or gemcitabine and paclitaxel (GEM/PTX) by scRNAseq. We found striking similarity in the pathways enriched in TAMs from mice treated with GEM/PTX compared to vehicle and pathways found when we stratified TAMs in vehicle treatment mice by p21 expression (Fig. 4 P). Similarly, TAMs from GEM/PTX treated KPC mice showed higher expression of the p21^High^ gene signature when compared to vehicle. (Fig. S2 J). These data suggested that p21 was induced by both stromal interaction and amplified by chemotherapy treatment, and correlated with inflammatory and likely immunosuppressive phenotypes in PDAC TAMs. Next, we analyzed the p21^High^ TAM signature in TCGA data sets and found strong correlation with signatures of “T cell exhaustion” (Tirosh et al., 2016) and “immune escape” (Lin et al., 2007) (Fig. 4 Q). Additionally, the p21 signature strongly correlated with CSF1 expression (Fig. 4 Q). These data suggest that stromal-CSF1 induced p21 expression in TAMs may drive dysfunctional T cell mediated tumor control.

### Expression of p21 drove the tumor promoting phenotype in macrophages

To better understand the impact of induction of p21 expression on the macrophage phenotype, and on the PDAC TME, we engineered a mouse designed to constitutively express p21 in myeloid cells. The construct contained the *p21* gene under the control of a CAG promoter and a lox-stop-lox case. Downstream of the *p21* gene, the construct also contained an internal ribosome entry site (IRES) and YFP gene for visualization. The construct was then integrated into the ROSA locus of pure C57/B6 mice (ROSA-CAG-LSL-p21-IRES-YFP, p21^+/wt^) (Fig. 5 A). Then, p21^+/wt^ mice were crossed with LysMCre mice to specifically induce p21 expression in macrophages. The resulting LysM^+/+^/p21^+/wt^ mice were termed “p21 constitutive expression” (p21^CE^) mice.

**Figure 5.**
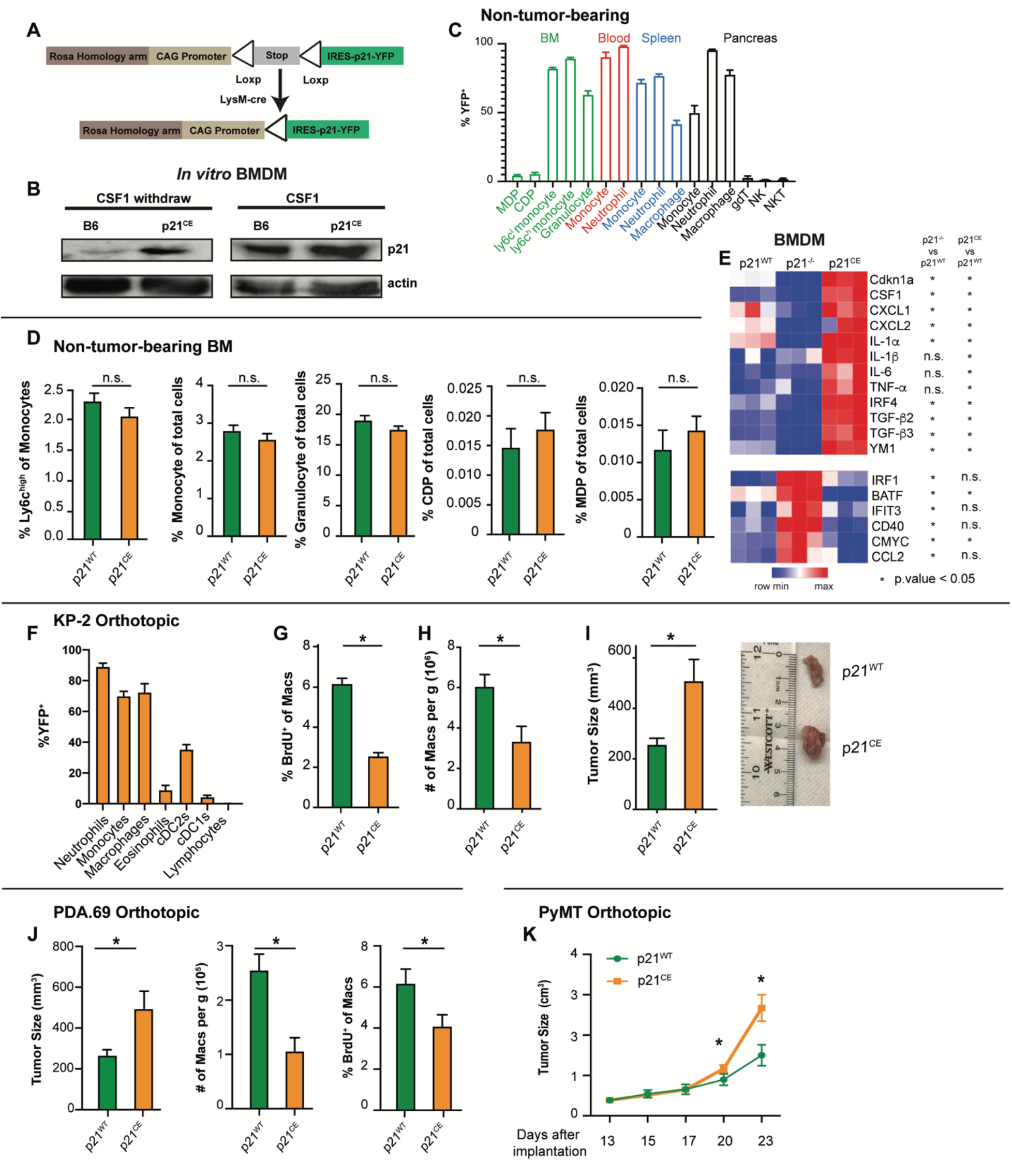
Expression of p21 drives tumor promoting phenotypes in macrophages. **(A)** Genetic loci for the p21^CE^ model. **(B)** Immunoblot for p21 expression in p21^CE^ or B6-derived bone marrow-derived macrophages (BMDMs) with or without 10ng/ml of colony stimulating factor-1 (CSF1) treatment for 24 h. Experiments were consistent in two independent repeats. **(C)** Bar plot displaying the percentage of YFP^+^ cells in non-tumor-bearing p21^CE^ mice; n = 4. **(D)** Bar plot showing flow cytometry quantification of cellular composition in non-tumor-bearing bone marrow from p21^CE^ and p21^WT^ mice; n = 6–9 mice/group. **(E)** Heat map displaying gene expression analysis of BMDMs derived from non-tumor-bearing p21^WT^, p21^−/−^, and p21^CE^ mice treated with 10ng/ml of CSF1 for 24 h, by RT-qPCR; n = 3/group, data was consistent from three independent repeats. **(F)** Flow cytometry quantification of YFP^+^ cells in p21^CE^ mice bearing orthotopic KP-2 tumors; n = 6–7 mice. **(G,H)** Quantification of BrdU^+^ macrophages and density of macrophages in tumors of p21^CE^ and p21^WT^ mice; n = 6–7 mice/group. Data were pooled across multiple independent experiments. **(I)** Bar plot displaying the tumor sizes in p21^CE^ and p21^WT^ mice, 21–27 days following orthotopic implantation of KP-2 tumor cells; n = 8–10 mice/group. **(J)** Bar plot displaying tumor sizes, density of macrophages, and quantification of BrdU^+^ macrophages from p21^CE^ and p21^WT^ mice, 21–23 days after the orthotopic implantation of the PDA.69 cell line; n = 8–10 mice/group. Data were pooled from multiple independent experiments. **(K)** Caliper measurement of orthotopic PyMT in p21^WT^ and p21^CE^ mice; n = 6–8 mice /group. All graphs are expressed as the mean ± SEM. n.s., not significant; *p < 0.05. All *in vitro* assays were consistent across more than two dependent repeats. For comparisons between any two groups, Student’s two-tailed *t*-test was used.

To confirm that p21 expression was induced in macrophages from p21^CE^ mice, we measured p21 protein levels in BMDMs from p21^CE^ mice in the presence and absence of CSF1. We found BMDMs from p21^CE^ mice expressed significantly higher levels of p21 protein in the absence of CSF1 compared to control BMDMs (Fig. 5 B). However, in the presence of CSF1, which strongly induced p21 expression in wildtype BMDMs (Fig. 4, A and B), both p21^CE^ and p21^WT^ BMDMs had similar p21 expressions. These data indicated that macrophages from the p21^CE^ mouse model retained high p21 expression without stimuli and that the expression was at a physiological level comparable to CSF1 exposure or fibroblast co-cultures.

Given LysMCre is known to be expressed in various myeloid compartments, including granulocytes and monocytes (Abram et al., 2014), we next examined whether the hematopoietic system was altered in p21^CE^ mice. Flow cytometry analysis of non-tumor-bearing p21^CE^ mice revealed that YFP, a surrogate for transgenic p21, was mainly expressed in mature monocytes, macrophages, and granulocytes/neutrophils in the blood, bone marrow, spleen, and pancreas, but minimally expressed in bone marrow progenitors and lymphocytes (Fig. 5 C). Corresponding to the lack of expression in progenitor cells, we did not find major changes in the cellular composition of bone marrow or blood in p21^CE^ mice compared to controls, as assessed by flow cytometry or by complete blood count analysis (Fig. 5 D; Fig. S3, A, B and C). Taken together, these data suggested that p21 was expressed mainly in mature myeloid cells in p21^CE^ mice, but minimal in progenitors and it did not greatly impact hematopoiesis.

As shown above in the scRNAseq data and gene profiling analysis after p21 siRNA knockdown, p21 expression regulated the macrophage phenotype. To assess whether macrophages from p21^CE^ mice had similar phenotypic changes, we profiled gene expressions of BMDMs from p21^WT^, p21^CE^, and p21^−/−^ (Jax mice) mice in the presence of CSF1. We found that inflammatory cytokines/chemokines, CXCL1, CXCL2, IL-1*α*, IL-1*β*, IL-6, and TNF-*α* were upregulated in p21^CE^ mice but reduced or not changed in p21^−/−^ mice (Fig. 5 E). In addition, the interferon regulatory factor 4 (IRF4)-mediated macrophage alternative activated genes, YM1 and transforming growth factor beta (TGF-*β*), were also upregulated. In contrast, p21^−/−^ BMDMs had elevated levels of the interferon-related genes, IRF1, BATF, IFIT3 and CD40, which were consistent with the changes in macrophages with siRNA-mediated knockdown of p21 (Fig. 5 E and Fig. 4 H). Taken together, these data suggest that constitutive p21 expression regulates the macrophage phenotype and represses anti-tumor immunity.

Next, we examined the impact of constitutive p21 expression in myeloid cells on PDAC progression. We orthotopically implanted KP-2 cells into p21^CE^ and p21^WT^ mice and analyzed tumors at the end point by flow cytometry. Similar to YFP expression patterns in non-tumor-bearing mice, we found in PDAC tissues that the majority of TAMs, monocytes, and neutrophils were YFP^+^, but the vast majority of tumor infiltrating cDCs, lymphocytes, and bone marrow progenitors were YFP^−^ (Fig. 5 F). Corresponding to lack of expression in DCs, we found no major changes in the numbers of cDC1s and cDC2s in pancreatic tissues from p21^CE^ tumor-bearing mice (Fig. S3 D). Additionally, the number of other myeloid cells that were not largely dependent on proliferation was also not changed in p21^CE^ when compared to p21^WT^ (Fig. S3, D - F). With constitutive expression of p21, we found a reduction in TAM proliferation, as measured by BrdU, as well as a decrease in total TAM numbers (Fig. 5, G and H). These data suggest that local proliferation of TAMs is necessary to sustain a local TAM pool. Interestingly, while TAM depletion in other studies typically slowed tumor growth (Zhu et al., 2014; Borgoni et al., 2018; Candido et al., 2018), we saw a significant increase in tumor burden in p21^CE^ mice (Fig. 5 I). These data suggest that changes in myeloid phenotype mediated by p21 drives tumor progression. Before evaluating the phenotypic changes of TAMs in p21^CE^ mice, we examined the tumor promoting effects on other tumor models. Similar to orthotopic KP-2, the PDA.69 PDAC model (Lee et al., 2016) and PyMT mammary tumor model showed decreased TAM proliferations and numbers, but accelerated tumor progression (Fig. 5, J and K). Together, these data suggest that constitutive expression of p21 in myeloid cells reduces TAM proliferations and numbers, but also alters TAM phenotype to drive tumor progression.

### The p21 expression in macrophages led to an inflammatory but immunosuppressive phenotype

We next sought to explore how high p21 expression in myeloid cells affected their phenotype *in vivo*. We conducted scRNAseq analyses on sorted CD45^+^ cells from PDAC tissues in p21^WT^ and p21^CE^ mice. An unsupervised clustering algorithm identified 19 clusters (Fig. S4 A), which mainly included C1qa-expressing macrophages, Ly6C2-expressing monocytes, S100a8-expressing granulocytes, Cd3d-expressing T cells, and Ms4a1-expressing B cells (Fig. 6 A; Fig. S4 B). To assess transgene expression, we analyzed the expression of YFP sequences. Consistent with flow cytometry data, myeloid compartments, including macrophages, monocytes, neutrophils, and eosinophils had high YFP expressions, while DCs had minimal and non-myeloid cells had no expression (Fig. 6 B).

**Figure 6.**
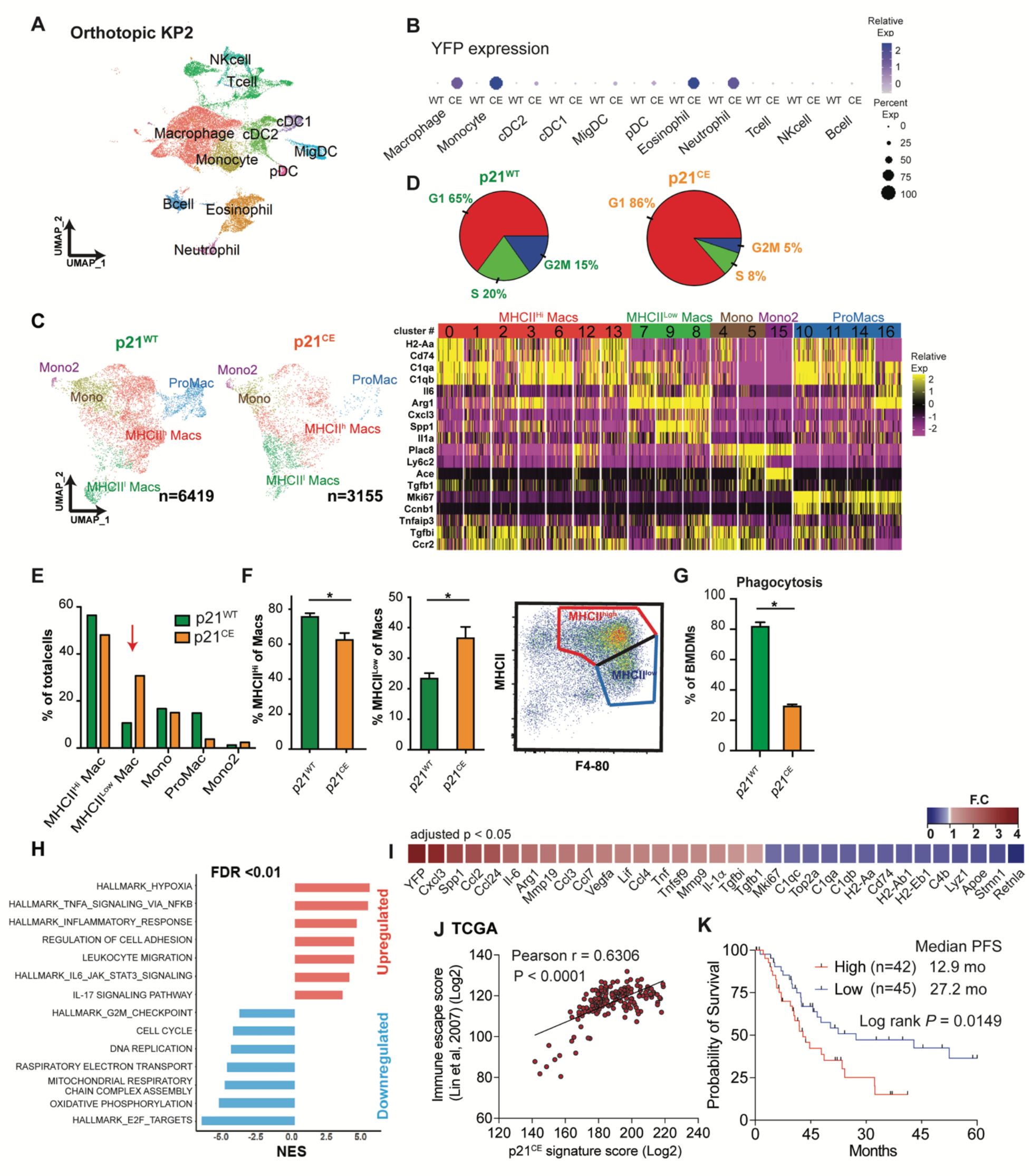
The p21 expression in macrophages led to an inflammatory but immunosuppressive phenotype. **(A)** UMAP dimensionality reduction plot of total CD45^+^ cells from p21^WT^ and p21^CE^ mice bearing orthotopic KP-2 tumors. Cells in each genotype were pooled from three mice and created as two libraries. Clusters were annotated with corresponding cell types. **(B)** Dot plot displaying YFP expression in each cell type between the two groups. The legend shows the dot size and corresponding percentage that are expressed as a color gradient of normalized expressions. **(C)** Reclustered UMAP plot of macrophage and monocyte clusters in **A** without cell cycle regression and split into p21^WT^ and p21^CE^, and annotated with major subpopulations. on the right, heat map showing key gene expressions in each subpopulation in **C**. **(D)** Pie chart showing cell cycle analysis of macrophages (MHCII^hi^, MHCII^low^, and ProMac) in tumors from p21^WT^ and p21^CE^ mice. **(E),** Bar plot showing quantification of each population between p21^WT^ and p21^CE^ mice identified in **C**. **(F)** Quantification of flow cytometry analysis of the percentages of MHCII^hi^ and MHCII^low^ macrophages from p21^CE^ and p21^WT^ mice bearing orthotopic KP-2 tumors with the representative gating strategy; n = 6–10 mice/group. Data were consistent in four independent repeats. **(G)** Barplot displaying quantification of fluorescent-bead^+^ bone marrow-derived macrophages from p21^WT^ and p21^CE^ mice. Data were consistent in three independent repeats. **(H)** Bar plot displaying Gene Set Enrichment Analysis results of comparing tumor-associated macrophages (TAMs) from p21^CE^ to p21^WT^ mice. The key upregulated and downregulated pathways are shown with a false discovery rate < 0.01. **(I)** Heat map showing the key differentially-expressed genes (DEGs) comparing TAMs from p21^CE^ and p21^WT^ mice. DEGs were filtered with an adjusted p < 0.05 and fold-change > 1.3 or < 0.75. All gene expressions were normalized by SCTransform. **(J)** Correlation plots with Pearson coefficients (r) of p21^CE^ signature score (included genes with LogFC >0.75) vs. Immune escape score from TCGA PDAC PanCancer Atlas study (n=180). **(K)** Kaplan-Meier survival analysis of PDA patients from TCGA whose samples were stratified by expression of the p21^CE^ signature (LogFC >0.75) by quartiles. All graphs are expressed as the mean ± SEM. n.s., not significant; *p < 0.05 using the *t*-test, except for **I** where the Bonferroni-corrected adjusted p-value was used.

To more accurately define myeloid subpopulations identified by scRNAseq and evaluate the phenotypic changes in each, starting from TAMs, we computationally separated macrophage/monocyte clusters and reanalyzed these at a higher resolution. This approach generated 17 clusters, which were grouped into four major populations, including macrophages with high MHCII expression (MHCII^hi^ Macs), low MHCII expression (MHCII^low^ Macs), monocytes (Mono, Mono2), and proliferating macrophages (ProMacs) (Fig. 6 C). After identifying major macrophage subsets, we first performed cell cycle analysis on all macrophages and confirmed that their proliferations were reduced (Fig. 6 D). Second, we observed that a higher percentage of TAMs in p21^CE^ was in the MHCIl^low^ cluster, and that this change was also observed at the protein level by flow cytometry (Fig. 6, E and F), indicating that TAMs in p21^CE^ potentially had impaired cross-presentation. Third, we performed Gene Set Enrichment Analysis (GSEA) between p21^CE^ TAMs and p21^WT^ TAMs and found that consistent with *in vitro* experiments, TAMs in p21^CE^ were enriched in TNF-*α* signaling, as well as pathways associated with hypoxia and inflammatory responses (Fig. 6H; Fig. S5 A). Notably, we also observed downregulation of genes associated with antigen processing and presentation of H2-Aa, H2-Ab1, H2-Eb1, and Cd74, and with the complement components of C1qa, C1qb, and Lyz, whereas tissue remodeling markers of Arg1, Mmp19, Vegfa, and Mmp9 were upregulated in TAMs from p21^CE^ tumor-bearing mice (> 1.5-fold, adjusted p < 0.05) (Fig. 6 I). Taken together, these data suggest that TAMs in p21^CE^ are more inflammatory, characterized by high TNF-*α*signaling, and are more immunosuppressive, characterized by both impaired anti-tumor functions and expressions of M2-like gene signatures. In addition, we found an increase of eosinophils within the TME of PDAC from p21^CE^ tumor-bearing mice (Fig. S4 C), which further illustrated that the TME was more inflammatory.

To further confirm that the p21^CE^ model recapitulated the characteristics of p21^High^ TAMs identified in mouse PDAC tissues in Fig. 4 K, we examined the expression levels of p21^High^ gene signature defined in Fig. 4 M in TAMs from p21^CE^ and p21^WT^ tumor-bearing mice. We found that TAMs in p21^CE^ expressed significantly higher levels of the p21^High^ gene signatures (Fig. S4 D). In addition, a gene encoded for the common *γ* chain of the FC receptor (*Fcer1g*) was significantly reduced in p21^High^ TAMs across three mouse scRNAseq datasets in Fig. 4 M. Cross-linking of Fc*γ*Rs and the common *γ* chain is required for IgG-mediated response and phagocytosis (Castro-Dopico and Clatworthy, 2019). Therefore, we evaluated whether p21^CE^ macrophages had impaired Fc*γ*R-mediated phagocytosis. We cultured BMDMs from p21^CE^ or p21^WT^ non-tumor-bearing mice with IgG-coated beads and found significantly less phagocytosis in p21^CE^ BMDMs (Fig. 6 G). These data suggest TAMs with high p21 expression have impaired effector functions which could contribute to tumor progression. Finally we analyzed a gene expression signature derived from TAMs in p21^CE^ mice in human PDAC expression datasets. Our analysis found that the p21^CE^ signature was also associated with “immune escape” signatures (Lin et al., 2007) and poor progression free survival (Fig. 6, J and K).

To understand the changes in other myeloid cells from p21^CE^ mice, we compared the numbers of significantly changed genes in each myeloid population between the two genotypes. We found that TAMs showed the largest number of differentially expressed genes (DEGs) (80 genes), followed by monocytes (34 genes), and only a few genes in neutrophils and granulocytes (Fig. S4 F). These data suggest macrophages are likely the predominant driver of tumor burden differences. To confirm macrophage contribution to the tumor difference between the two genotypes, we administered *α*CSF1 IgG and clodronate-containing liposomes to p21^CE^ and p21^WT^ tumor-bearing mice throughout tumor development. We found that the number of TAMs was significantly reduced, while the number of monocytes did not after the treatment in both genotypes of mice (Fig. S4, I and J). Only in the setting of macrophage depletion were the tumor promoting effects observed in p21^CE^ mice abolished (Fig. S4 H). Therefore, these data suggest that macrophages are the main driver for tumor acceleration in p21^CE^ mice.

Although YFP was not significantly expressed by DCs, DCs play a critical role in antigen processing and presentation as well as CD8^+^ T cell activity and could potentially affect tumor progression (Gardner and Ruffell, 2016). To evaluate the changes in DCs in p21^CE^ tumors, we reclustered DC populations from scRNAseq data at a higher resolution and identified seven major subsets: cDC1, cDC2a, cDC2b, migratory DC (MigDC), pDC, and proliferating cDC1 and cDC2 (Fig. S4 E). The cDC1 expressed classical DC1 markers of *Xcr1*, *Clec9a*, and also *Baft3* and *Irf8*, while the cDC2 subsets expressed *Cd11b*, *Irf4*, and *Sirpa*, and were further separated into cDC2a and cDC2b based on *Epcam* expression (Merad et al., 2013; Kaplan, 2017). We did not observe significant changes in the percentages of cDC1s, cDC2s, migratory DCs, and proliferating DCs as the total number of DCs between two genotypes, nor did we observe a change in genes associated with cross-presentation. We saw a decrease in pDCs and an increase of cDC2bs as the percentage of total DCs (Fig. S4 F). Because pDCs are one of the major producers of type-I interferon (Koucký et al., 2019) and could potentially drive anti-tumor immunity, this reduction could impact tumor immune suppression.

### The p21 expression in macrophages impaired effector T cells

To determine if impaired antigen processing and presentation in macrophages directly affected T cell numbers and functions, we reanalyzed T cell clusters from the scRNAseq experiment at a higher resolution. Unsupervised clustering generated 12 clusters and were manually assigned into natural killer cells (NK cells), regulatory T cells (T_Regs_), two clusters of CD4^+^ (CD4#1 and CD4#2), two clusters of CD8^+^ (CD8#1 and CD8#2), double negative T cells (DNs), and a gamma delta T cell based on known cell type markers (Fig. 7 A). Among CD8^+^ T cells, cluster #2 expressed the higher effector genes, *Gzma*, *Gzmb*, and *Cd74*, and therefore was considered as cytotoxic effectors (Fig. 7 A). We observed that this CD8^+^ effector cluster was reduced as a percentage in p21^CE^ tumor-bearing mice (Fig. 7 B) and the expressions of effector genes, *Gzma*, *Gzmk*, *Klrg1*, were also significantly lower (Fig. 7 D). In contrast, we saw an increase in the percentage of CD4#2 T cell populations, which are T_H_2 polarized, with high levels of Gata3, *IL-4* and *IL-13* (Fig. 7 B) (Zheng and Flavell, 1997). If mapping the upregulated genes in cytotoxic CD8^+^ T cells from p21^CE^ tumors to known signaling pathways, we found enrichment in apoptosis and IL-2-STAT5 signaling, suggesting overexpressed p21 in macrophages may cause more cytotoxic CD8^+^ T cell death (Fig. 7 C). To confirm this, we co-cultured activated CD8^+^ T cells with BMDMs generated from p21^CE^ and p21^WT^ mice *in vitro*, and found p21^CE^ BMDMs led to more apoptosis of CD8^+^ T cells, measured by 7-AAD (Fig. 7 E). To extend the findings to human PDAC patients, we analyzed the correlations between the p21^CE^ signature in TAMs with “T cell exhaustion”(Tirosh et al., 2016) and found strong positive correlations (Fig. 7 K). Taken together, these data suggest high p21 expression in TAMs dampens cytotoxic CD8^+^ T cell mediated tumor control.

**Figure 7.**
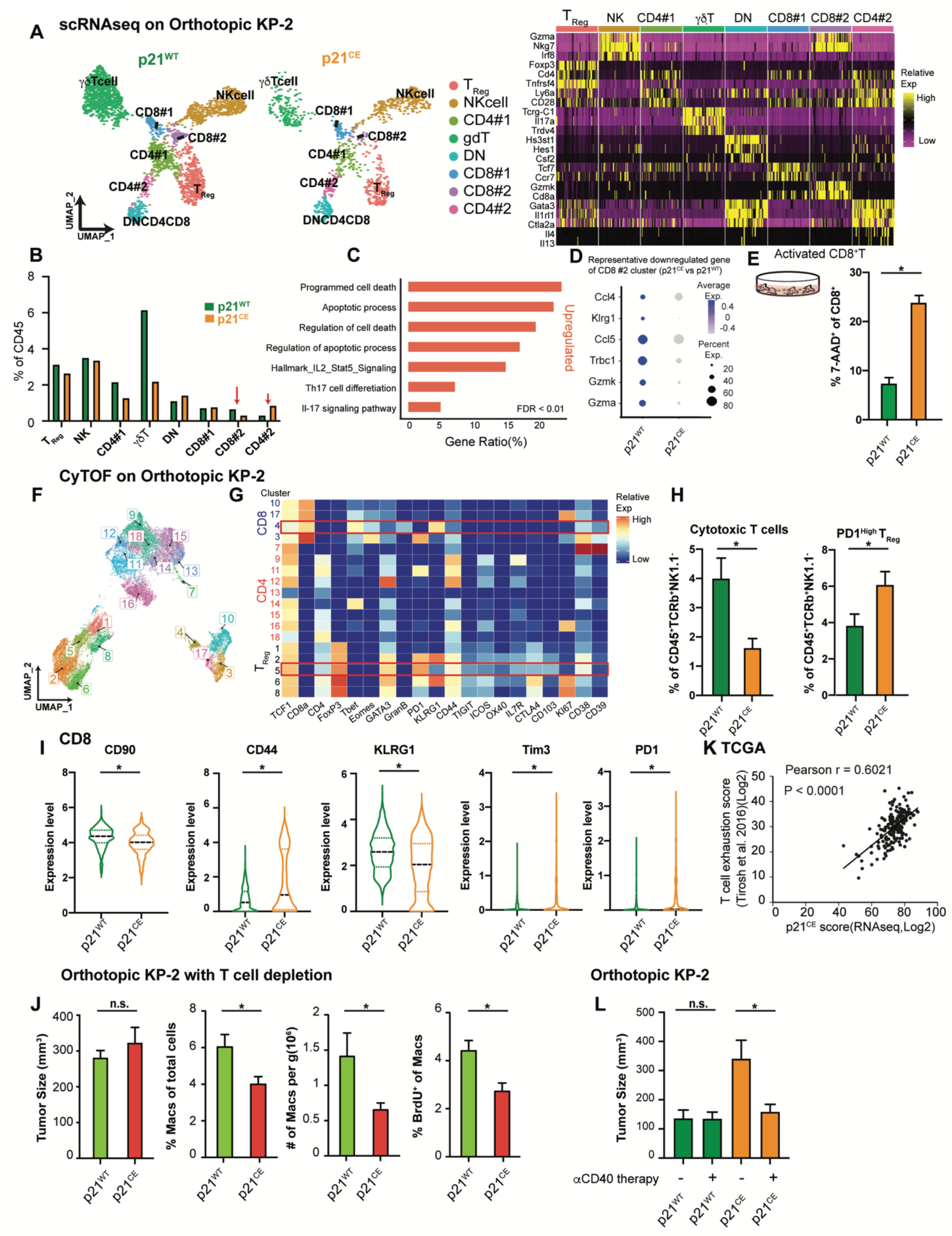
The p21 expression in macrophages impaired effector T cells. **(A)** UMAP dimensionality reduction plot of selected lymphocytes (clusters 6, 7, 8, 13, and 17 in Fig. S4, A and B) from p21^WT^ and p21^CE^ orthotopic KP-2 tumors. Clusters were annotated with corresponding cell types and heat maps displaying selected gene expressions in each cell type. **(B)** Bar graph displaying the composition of each cell type as the percentage of total CD45^+^ cells in p21^WT^ and p21^CE^ tumor-bearing mice. CD8#2 and CD4#2 are highlighted with red arrows. **(C)** Bar graph displaying the upregulated pathways in the CD8#2 cluster from p21^CE^ using overrepresentation analysis of differentially-expressed genes (DEGs) to known biological functions (Gene Ontology, Kyoto Encyclopedia of Genes and Genomes, REACTOME, and the Molecular Signal Database). DEGs were filtered with a value of p < 0.05, fold-change > 1.2, and past MAST test. **(D)** Table showing the differentially expressed genes comparing CD8#2 cluster from p21^CE^ to p21^WT^ with p.value < 0.05. **(E)** Bar plot displaying the percentage of 7-AAD^+^CD8^+^ T cells activated with CD3/CD28 Dynabeads (Gibco) when cocultured with BMDMs from p21^CE^ and p21^WT^ mice for 48 h. Data were consistent in three independent repeats. **(F)** UMAP plot of selected CD45^+^TCRb^+^CD90^+^NK1.1^−^TCR^−^*γδ*T^−^ cells from p21^CE^ and p21^WT^ orthotopic KP-2 tumors with clusters annotated; n = 7 mice/group. **(G)** Heat map displaying the feature expressions in each cluster. Cytotoxic T cells (cluster 4) and PD1^High^ T_reg_ (cluster 5) were highlighted. **(H)** Bar plot showing the percentages of cytotoxic T cells and PD1^High^ T_reg_ in p21^WT^ and p21^CE^ tumors. **(I)** Violin plot visualizing the expression levels of CD90, CD44, KLRG1, TIM3, and PD1 in the CD8 cluster between tumors from two genotypes. **(J)** Bar graphs showing the tumor burden, macrophages as the percentage of total cells, or as per gram of tissue, and the percentage of BrdU^+^ macrophages between p21^WT^ and p21^CE^ orthotopic KP-2 tumors after *α*CD4/CD8 treatment; n = 6 mice/group. **(K)** Correlation plots with Pearson coefficients (r) of p21^CE^ score vs. T cell exhaustion score from TCGA PDAC PanCancer Atlas study (n=180). **(L)** Bar graph showing the tumor burdens of p21^WT^ and p21^CE^ mice bearing orthotopic KP-2 tumors with or without CD40 agonist and gemcitabine treatment. n= 5-6 mice/group. All graphs are expressed as the mean ± SEM. n.s., not significant; *p < 0.05 for comparisons between two groups **E,H,J,L**, Student’s two-tailed *t*-test was used. For comparisons in **I**, the Bonferroni-corrected p-value was used.

To corroborate these findings, we used a T cell-focused CyTOF panel. CD45^+^TCRb^+^CD90^+^NK1.1^−^ TCR^−^*γδ*T^−^ cells were selected for further clustering based on 20 T cell functional markers. This approach generated 18 clusters that could be mainly grouped into three major populations: CD4^+^ T cells, regulatory CD4^+^ T cells (T_Regs_), and CD8^+^ T cells (Fig. 7, F and G). We next evaluated changes in each subpopulation and found a significant decrease in the numbers of cytotoxic effectors (cluster 4), which expressed high levels of granzyme B and KLRG1. In addition, we observed an expansion of the CD4^+^T_reg_ (cluster 5) that expressed high levels of PD1 (Fig. 7 H). In addition, we found that CD8^+^ T cells as a whole in p21^CE^ tumors expressed lower levels of KLRG1 and CD90, but higher levels of CD44, Tim3, and PD1, indicating a more exhausted and less functional phenotype (Fig. 7 I). Finally, to determine whether accelerated tumor progression in p21^CE^ mice was driven by T cells, we depleted CD4^+^ and CD8^+^ T cells in both p21^CE^ and p21^WT^ mice through injection of *α*CD4 IgG and *α*CD8 IgG. We no longer observed difference in tumor burdens between the two groups (Fig. 7 J). These data suggest that p21-driven TAM immunosuppressive phenotype not only reduces the number of anti-tumor T cells but also impairs the functions of remaining T cells.

We next asked whether innate immune agonist therapy, CD40 agonist, could reeducate TAMs and restore their effector functions (Coveler et al., 2020). To test this, we treated p21^CE^ and p21^WT^ mice bearing orthotopic KP-2 tumors with CD40 agonist therapy and found that while the dual treatment had limited effect on p21^WT^ mice, it dramatically reduced the tumor burden in p21^CE^ mice (Fig. 7 L). These data suggest, although stromal or chemo-induced p21 expression drives an inflammatory and immunosuppressive phenotype in TAMs, these same pathways may make tumor uniquely susceptible to CD40 agonist therapy.

## Discussion

Macrophage proliferation has been observed in several non-cancer pathological conditions, including helminth infections (Jenkins et al., 2011), atherosclerosis (Tang et al., 2015), and obesity-associated adipose tissues (Amano et al., 2014). In these conditions, proliferation of macrophages, albeit under the control of different factors, is necessary to sustain total macrophage numbers at each tissue site. In our studies, we found in pancreatic tumors that macrophage proliferation was mainly driven by CAF-derived CSF1. These data implied that although the general need for macrophage expansion was common, the activated signaling pathways and resulting macrophage phenotypes were largely tissue- and context-dependent. Stromal rich tumors may increase TAM numbers more frequently by local proliferation. Interestingly, CSF1 levels were reported to be higher in the blood of patients suffering from melanoma, breast cancer, or pancreatic cancer. In these patients and also in corresponding mouse models, macrophages were found to be proliferative (Bottazzi et al., 1990; Franklin et al., 2014; Tymoszuk et al., 2014). These data suggested that CSF1-driven macrophage proliferation was common in multiple cancer types.

An earlier study examined the CSF1 effects on CSF1R-expressing human breast cancer cell lines, and found that CSF1 inhibited cell proliferation through inducing p53 independent, but MAPK-dependent, p21 expression (Lee et al., 1999). This result may seem contradictory to ours as we showed CSF1 induced BMDM proliferation. However, we also showed that knocking-down p21 expression or constitutively expressing it promoted or inhibited macrophage proliferation. These data suggested that CSF1 induction of p21 in macrophages acted as a checkpoint for S phase entry. The ultimate cell cycle transit required additional signaling, and the signals could be synthesized according to the expression level of p21. One group reported that Raf signal intensity determined either induction of DNA synthesis or inhibition of proliferation in fibroblasts by p21^Cip1^ expression levels (Sewing et al., 1997). A recent study further showed that p21 not only determined the cell cycle fate of mother cells but could also be carried into daughter cells and regulated the proliferation after mitosis (Yang et al., 2017). Therefore, it is not surprising that the p21 expression level is known to protect cells from chemotherapy-induced apoptosis (Hsu et al., 2019).

Aside from p21’s canonical role as a cell cycle checkpoint, several groups reported its role in regulating inflammation, with some contradictory results. One group demonstrated that p21^−/−^ mice were more sensitive to LPS-induced septic shock due to inflammation (Trakala et al., 2009). Likewise, p21^−/−^ mice showed enhanced experimental inflammatory arthritis and severe articular destruction (Mavers et al., 2012).Contrastingly, in a serum transfer model of arthritis, p21^−/−^ mice were more resistant (Scatizzi et al., 2006). Furthermore, disruption of p21 attenuated lung inflammation in mice (Yao et al., 2008). These data suggested that regardless of whether p21 promoted or inhibited inflammation, it was established that p21 regulated inflammation. In a chronic pancreatitis model, one study found that p21 expression was significantly increased overall, while knocking-down its expression resolved inflammation and prevented pancreatic injury through reducing the release of NF-*κ*B-mediated proinflammatory cytokines, such as TNF-*α*, IL-6, and CXCL1(Seleznik et al., 2018). These data suggested that at least in the pancreas, p21 played a role in promoting inflammation, independent of KRAS mutations that are commonly observed in PDAC and are known to drive inflammation (Kitajima et al., 2016). However, this study did not identify the main drivers for p21-mediated inflammation.

Macrophages are known to exhibit plasticity, which gives them the capability to quickly respond to environmental challenges. Expression levels of p21 could be an important regulator in macrophage plasticity. Expression of p21 inhibited macrophage activation during LPS-induced septic shock, as p21^−/−^ macrophage expressed higher levels of CD40 and enhanced activation of NF-*κ*B (Trakala et al., 2009). One study further demonstrated that expression of p21 acted more like a buffer system for inflammation as it could adjust the equilibrium between p65-p50 and p50-p50 NF-*κ*B pathways to mediate macrophage plasticity in LPS treatment (Rackov et al.). However, none of these studies investigated p21 effects on macrophage polarization in tumor settings. From scRNAseq data, we showed that stratifying macrophages based on p21 expressions into p21^Hi^ and p21^Low^ resulted in two phenotypically distinct macrophages independent of the cell cycle, with the first being more inflammatory. TNF-*α* and NF-*κ*B were upregulated when p21 expression was high, which is consistent with previous findings. We further illustrated that constitutive expression of p21 in macrophages impaired their phagocytosis capabilities *in vitro*, lowered expression of genes associated with antigen cross-presentation in orthotopic PDAC tumors, and hindered cytotoxic T cell functions, which eventually led to faster tumor progression. These observations are important because as we showed both stromal interaction and therapeutic interventions targeting cell cycle could induce p21 expression in TAMs and lead to an inflammatory yet immunosuppressive phenotype. Given TAMs are usually abundant in TME, these p21-driven phenotypic changes could eventually lead to resistance for treatments.

We also found that in human and mouse PDACs, although p21 expression was highest in macrophages, it was expressed by other myeloid populations. If p21 regulates inflammatory responses through NF-*κ*B in macrophages, it is possible that other immune cells mediate inflammation, like granulocytes and neutrophils, which could also be polarized by p21 in a similar way. One group observed that p21 expression in neutrophils regulated inflammation in infections (Martin et al., 2016). In addition, we observed that p21 expression was induced by chemotherapy not only in macrophages, but also in other myeloid cells, which suggested that inflammatory but immunosuppressive phenotypes could be further strengthened by myeloid cells, in addition to macrophages.

Understanding how the TME and cancer cell intrinsic factors regulate macrophage tumor supportive vs. tumor suppressive functions is critical to therapeutically targeting TAMs in cancer patients. In total, our data suggested that CAF-induced macrophage proliferation was important for sustaining TAM number and induction of p21, which also resulted in immunosuppression and tumor progression. Lastly, expression of p21 in TAMs might sensitize tumors to CD40 agonist treatment.

## Materials and methods

### Contacts for reagent and resource sharing

Further information and requests for resources and reagents should be directed to and will be fulfilled by the lead contact, David G. DeNardo.

### Murine PDAC models

Mice were maintained in the Laboratory for Animal Care barrier facility at the Washington University School of Medicine. All studies were approved by the Washington University School of Medicine Institutional Animal Studies Committee.

KPC mice (p48-Cre;Kras^LSL-G12D^;Trp53^fl/fl^) used in these studies have been rapidly bred to the C57Bl/6J background in our laboratory using speed-congenics and further backcrossed more than five times. All mice were housed, bred, and maintained under specific pathogen-free conditions in accordance with NIH-AALAC standards and were consistent with the Washington University School of Medicine IACUC regulations (protocols #20160265 and #19-0856).

The KP-1 cell line was derived from PDAC tissues of the 2.2-month-old p48-CRE^+^/LSL-Kras^G12D^/p53^flox/flox^ (KPC); the KP-2 cell line was derived from the 6-month-old p48-CRE^+^/LSL-Kras^G12D^/p53^flox/+^ mice(KP^fl/+C^) (Jiang et al., 2016). The KI cell line was derived from the Pdx1-Cre;LSL-Kras^G12D^;Ink/Arf^fl/fl^ as previously described (Mitchem et al., 2013). Cells were grown on collagen-coated tissue culture flasks for < 12 passages, and were tested for cytokeratin-19, smooth muscle actin, vimentin, and CD45 to verify their carcinoma identity and purity. The PDA.69 cell line was a kind gift from Dr. Gregory L. Beatty, and was maintained in tissue culture flasks with DMEM supplemented with 1% glutamax and 0.167% gentamycin for less than 13 passages. To establish orthotopic PDAC models, either 50,000 or 200,000 KP-2 cells, and 10,000 or 50,000 PDA.69 cells in 50 μL of Cultrex (Trevigen, Gaithersburg, MD, USA) were injected into the pancreas of 8–12-week-old C57BL/6 mice or transgenic mice according to published protocols (Kim et al., 2009). Tumor-bearing mice were sacrificed when the palpable tumor size was > 1 cm (21–27days).

### Other mouse models

The p21^CE^ mouse was developed at the Washington University Mouse Embryonic Stem Cell Core using the construct of Cdkn1a (p21, accession #NM_007669). Briefly, the construct contained the p21 gene under the control of a CAG promoter and a lox-stop-lox case. Downstream of the p21 gene, the construct also contained an internal ribosome entry site (IRES) and YFP gene for visualization. The construct was then integrated into the ROSA locus of pure C57/B6 mice (ROSA-CAG-LSL-p21-IRES-YFP) and injected into C57 blastocyst (p21^+/wt^). Successful chimeras were selected and verified by DNA sequencing across ROSA junctions (primers are listed in Table S2) and subsequent founder mice were identified via genomic PCR (primers are listed in Table S2). Then, p21^+/wt^ mice were crossed with LysMCre mice to specifically induce p21 expression in macrophages. The resulting LysM^+/+^/p21^+/wt^ mice are termed “p21 constitutive expression” (p21^CE^) mice.

### Tissue harvest

Mice were euthanized by intracardiac perfusion with 15 mL of phosphate-buffered saline (PBS)-heparin under isoflurane anesthesia. Blood was obtained by cardiac puncture and deposited in heparin-PBS (Alfa Aesar Lonza, Haverhill, MA, USA) solution. Blood was then incubated in red blood cell lysis buffer (Biolegend, San Diego, CA, USA) for 10 min on ice and quenched with 1% fetal bovine serum (FBS; Atlanta Biologicals, Flowery Branch, GA, USA) containing PBS. Normal and tumor tissues were manually minced and digested in 20 mL of Hank’s Balanced Salt Solution (Thermo Fisher Scientific, Waltham, MA, USA) supplemented with 2 mg/mL of collagenase A (Roche, Basel, Switzerland) and 1× DNase I (Sigma-Aldrich, St. Louis, MO, USA) for 30 min (20 min for normal tissue) at 37°C with agitation. After digestion, the cell suspensions were quenched with 5 mL of PBS and filtered through 40 μm nylon mesh. The filtered suspensions were then pelleted by centrifugation (1,800 rpm for 4 min at 4°C) and resuspended in flow cytometry buffer [PBS containing 1% bovine serum albumin (BSA) and 5 mM EDTA] as a single cell suspension.

### Flow cytometry

Following tissue digestion, single cell suspensions were blocked with rat anti-mouse CD16/CD32 antibodies (eBioscience, Waltham, MA, USA) for 10 min on ice, and pelleted by centrifugation. The cells were subsequently labeled with 100 µL of fluorophore-conjugated anti-mouse extracellular antibodies at recommended dilutions for 30 min on ice in flow cytometry buffer. Intracellular staining was conducted using eBioscience Transcription Factor Staining Buffer using the manufacturer’s recommended procedures. All antibodies are listed in Table S3. For live analysis of YFP, fluorophore-labeled cells were analyzed immediately without fixation on X-20 cytometers.

For proliferation assays, mice were injected with BrdU, 1 mg i.p. at 3 h prior to sacrifice. A BD Biosciences Cytofix/Cytoperm kit (BD Biosciences, San Jose, CA, USA) was used following extracellular staining to stain for BrdU.

### Human samples

Human PDAC samples were obtained from consenting patients diagnosed at Washington University and the Siteman Cancer Center. Patients underwent pancreaticoduodenectomy. The Washington University Ethics committee approved the study under IRB protocol #201704078.

### Mass cytometry

Human tumor samples were collected on different days right after surgery and digested in Hank’s Balanced Salt Solution supplemented with 2 mg/mL collagenase A (Roche), 2.5 U/mL hyaluronidase (Sigma-Aldrich), and DNase I at 37°C for 30 min with agitation to generate single cell suspensions. Cell suspensions were counted and stained in 5 μM cisplatin per million cells for exactly 3 min on ice and washed with Cy-FACS buffer (PBS, 0.1% BSA, 0.02% NaN_3_, and 2 mM EDTA) twice. The cells were then incubated with FcR blocking reagent plus surface-antibody cocktail for 40 min on ice. After incubation, surface marker-stained cells were washed twice with Cy-FACS buffer. Cells were then fixed with 4% paraformaldehyde (PFA) for 10 min on ice and permeabilized with permeabilization buffer containing the intracellular stain cocktail (Invitrogen, Carlsbad, CA, USA) for 40 min. All antibodies are listed in Table S5. The cells were then washed and fixed a second time in 4% PFA in PBS at 4°C at least overnight. One day prior to acquisition, the cells were washed twice and stained with 200 μL of DNA intercalator per million cells. Cells were acquired on a CyTOF2 mass cytometer (South San Francisco, CA, USA) and were normalized with the MATLAB normalizer (v.7.14.0.739 run in MATLAB R2012a) (Finck et al., 2013). The normalized data were uploaded into Cytobank and manually gated to exclude normalization beads, cell debris, dead cells, doublets, and CD45^−^ cells. The filtered sample from each individual specimen was then exported and batch normalized by the date of acquisition using the R Cydar package NormalizeBatch function (mode = “range”) to compute a quantile function from the pooled distribution of the input expression data (Lun et al., 2017). In brief, batch expression was scaled between the upper and lower bounds of the pooled reference distribution, with zero values fixed at zero. A total of 10,245 events per batch of corrected sample was then visualized using the standard t-SNE algorithm in Cytobank. Populations of interest were manually gated and verified based on lineage marker expressions.

For mouse samples in Fig. 7 F-I, seven mice per group were individually stained for surface and intracellular stains (the antibodies are listed in Table S6), and fixed overnight as described above. Each sample was then barcoded with a unique combination of palladium metal barcodes using the manufacturer’s instructions (Fluidigm). Following bar coding, the cells were pooled together and incubated overnight in 2% PFA containing 40 nM iridium nucleic acid intercalator (Fluidigm). On the day of acquisition, the barcoded samples were washed and suspended in water containing 10% EQ Calibration Beads (Fluidigm) before acquisition on a CyTOF2 mass cytometer (Fluidigm). Sample barcodes were interpreted using a single cell debarcoder tool (Zunder et al., 2015). FCS files were then uploaded to Cytobank and manually gated to exclude normalization beads, cell debris, dead cells, and doublets. Classical T cells were classified as CD45^+^, Cisplatin^−^, Thy1.2^+^, NK1.1^−^, TCRgd^−^, and TCRb^+^. All T cells were exported as new FCS files and analyzed using the R CATALYST package (Nowicka et al., 2017) in R, version 3.8.2 (The R Project for Statistical Computing, Vienna, Austria). In brief, FCS files were down-sampled to equivalent cell counts, before clustering with the R implementation of the Phenograph algorithm (Levine et al., 2015). All markers were used for clustering analysis except markers used for T cell gating (see above). Dimensional reduction and visualization were performed using the UMAP algorithm (McInnes et al., 2020). Finally, differential cluster abundance testing was performed with the R diffcyt package, utilizing a generalized linear mixed model (Weber et al., 2019).

### Macrophage depletion

In Fig. 3 F, 8–12-week-old C57BL/6 mice were orthotopically implanted with 200,000 KP-2 cells. When the tumor was palpable, mice were intraperitoneally treated with one dose of 1 mg CSF1 neutralizing antibody (clone 5A1; BioXCell, Lebanon, NH, USA) and sacrificed at 12 and 24 h after treatments.

In Fig. S4, H - J, to deplete tissue resident macrophages, 8–12-weeks-old p21^CE^ and p21^WT^ mice were implanted orthotopically with 50,000 KP-2 cells on day 0, then were treated with three doses of CSF1 neutralizing antibody (1 mg, 0.5 mg, and 0.5 mg on days 3, 10, and 17) and two doses of clodronate-containing liposomes (200 µL each on days 5 and 12). Control mice were treated with the same doses/volumes of IgG (clone HRPN, BioXCell) and PBS liposomes.

### *In Vitro* co-culture and siRNA treatment

All cell lines were maintained in DMEM (Lonza, Basel, Switzerland) supplemented with 10% FBS (Atlanta Biological) and penicillin/streptomycin (Gibco, Gaithersburg, MD, USA). All cell lines tested negative for mycoplasma.

Pancreatic fibroblasts were harvested from the pancreas of healthy 8-week-old C57BL/6 mice, passaged three times on tissue culture plates, and tested negative for mycoplasma. An immortal pancreatic fibroblast cell line was established by passage more than 18 times. Soluble factors in primary pancreatic fibroblasts and immortal pancreatic fibroblasts medium were measured, compared, and found to be similar.

Bone marrow cells were obtained from both femur and tibia of the mouse and differentiated for five days in DMEM supplemented with 10ng of CSF1 (PeproTech, NJ, USA) for five days to generate BMDMs. A total of 75,000 fibroblasts or 50,000 KP-2 cells or both cell types were co-cultured with 100,000 BMDMs in 6-well cell culture plates (Costar, San Jose, CA, USA). BrdU was added 6 h prior to harvest at each time point. For Transwell assays, 150,000 fibroblasts were cultured in the Transwell assay with 200,000 BMDMs, and BrdU was added 6 h prior to harvest.

Small interfering RNAs (siRNAs) targeting mouse CSF1 and p21 were purchased from Integrated DNA Technologies (Coralville, IA, USA). Sequences are listed in Table S2. The siRNA transfections for primary BMDMs and pancreatic fibroblasts were performed using the Mouse Macrophage Nucleofector™ Kit (Lonza) and Nucleofector^™^ 2b Device (Lonza) with prewritten program Y-001 for BMDMs and V-013 for fibroblasts, following the manufacturer’s instructions. RNA and protein from transfected primary cells were harvest 24 h after the transfections.

### Microarray and RT-qPCR analysis

Total RNA was isolated from BMDMs derived from p21^CE^, p21^WT^,or p21^−/−^, or from siRNA targeting for p21-treated BMDMs using the E.N.Z.A. Total RNA Kit (Omega Chemicals, Cowpens, SC, USA) according to the manufacturer’s instructions. Microarrays were performed on p21 knocked-down BMDMs with the treatment of tumor-conditioned medium for 24 h. A differential gene list was generated with detected fold- changes > 1.5, adjusted p < 0.05. The filtered differential gene list was loaded into R and a hypergeometric test was used to compare known catalogs of functional annotations (enricher) with a FDR of p < 0.05. Top differentially-regulated genes are listed in Table S1. RNAs from BMDMs of p21^CE^, p21^WT^,and p21^−/−^ were reversed-transcribed to cDNAs by using the qScript cDNA SuperMix (QuantaBio, Beverly, MA, USA). Quantitative real-time PCR Taqman primer probe sets specific for targets listed in Table S7 (Applied Biosystems, Foster City, CA, USA) were used, and the relative gene expression for each target was determined on a ABI7900HT quantitative PCR machine (Applied Biosystems) using a Taqman Gene Expression Master Mix (Applied Biosystems). The threshold cycle method was used to determine fold-changes of gene expressions normalized to *Gapdh*, *Hprt,* and *Tbp*.

### ELISA and the cytokine array

Conditioned media from fibroblasts and tumor cells were harvested after changing the medium to 0.1% FBS for 24 h with > 80% confluency. The cytokine array were conducted by using a Proteome Profiler Mouse XL Cytokine Array kit (R&D Systems, Minneapolis, MN, USA) following the manufacturer’s instructions. The membranes from each conditioned medium were placed in an autoradiography film cassette and exposed to X-ray filming for 5–8 min. Positive signals were quantified by ImageJ software (National Institutes of Health, Bethesda, MD, USA). Conditioned media were concentrated using a Pierce Concentrator (Thermo Fisher Scientific) based on the manufacturer’s instructions. CSF1 levels were measured by a Mouse M-CSF Matched Antibody Pair Kit (ab218788) following the manufacturer’s instructions.

### Single cell RNA sequencing

Normal pancreas tissues were taken from three 10-week-old B6 mice, processed to single cell suspension as explained in the tissue harvest section, pooled together, and sorted for live macrophages (CD45^+^CD11b^+^F4/80^+^CD3^−^CD19^−^Siglecf^−^Ly6G^−^Ly6C^−^7AAD^−^) by using an Aria II cell sorter (BD Biosciences)

Pancreatic tumors were taken from three 1.5-month-old KPC mice, processed to a single cell suspension, pooled, and sorted for live macrophages and DC-enriched populations (CD45^+^CD3^−^CD19^−^ SiglecF^−^Ly6G^−^7AAD^−^).

Orthotopic KP-2 tumors were taken from p21^CE^ and p21^WT^ mice, and three from each genotype were pooled as one sample and sorted for live CD45+ cells (CD45^+^7AAD^−^). Two libraries were created for each genotype.

Sorted cells from each sample were encapsulated into droplets and libraries were prepared using Chromium Single Cell 3’v3 Reagent kits according to the manufacturer’s protocol (10x Genomics, Pleasanton, CA, USA). The generated libraries were sequenced by a NovaSeq 6000 sequencing system (Illumina, San Diego, CA, USA) to an average of 50,000 mean reads per cell. Cellranger mkfastq pipeline (10X Genomics) was used to demultiplex illumine base call files to FASTQ files. Files from the normal pancreas, pancreatic tumors, and orthotopic tumors were demultiplexed with > 97% valid barcodes, and > 94% q30 reads. YFP sequences were inserted into the mm10 reference (v.3.1.0; 10X Genomics) using the Cellranger Mkref pipeline. Afterwards, fastq files from each sample were processed with Cellranger counts and aligned to the mm10 reference (v.3.1.0, 10X Genomics) or mm10 containing YFP for p21^CE^ orthotopic tumor samples and the generated feature barcode matrix.

Human scRNAseq data were obtained from a publicly available dataset (Peng et al., 2019). FASTQ files were realigned to the human GRCh38 reference and generated feature barcode matrix, including 24 PDAC samples and 11 normal samples. However, only 21 PDAC samples and six normal samples successfully passed the Cellranger count function.

Mouse scRNAseq data (mPDAC GEMM-1) used in Fig. 3 A,B and Fig. 4 L,M and Fig. S2 F were obtained from a published paper (Hosein et al., 2019).

### Mouse scRNAseq data analysis

The filtered feature barcode matrix from the normal pancreas, KPC pancreatic tumors, and p21^WT^ orthotopic tumors were loaded into Seurat as Seurat objects (Seurat v.3). For each Seurat object, genes that were expressed in less than three cells and cells that expressed less than 1,000 or more than 8,000 genes, were excluded. Cells with greater than 6% mitochondrial RNA content were also excluded, resulting in 9,821 cells for normal, 6,091 for KPC tumors, and 16,904 for orthotopic tumors. SCTransform with default parameters was used on each individual sample to normalize and scale the expression matrix against the sequence depths and percentages of mitochondrial genes. Cell cycle scores and the corresponding cell cycle phase for each cell were calculated, and assigned after SCTransform based on the expression signatures for S and G2/M genes (CellCycleScoring). The differences between the S phase score and G2/M score were regressed-out by SCTransform on individual samples. Variable features were calculated for each sample independently and ranked, based on the number of samples they were independently identified (SelectIntegrationFeatures). The top 3,000 shared variable features were used for multi-set canonical correlation analysis to reduce dimensions and identify projection vectors that defined shared biological states among samples and maximized overall correlations across datasets. Mutual nearest neighbors (MNNS; pairs of cells, with one from each dataset) were calculated and identified as “anchors” (FindIntegrationAnchors). Multiple datasets were then integrated based on these calculated “anchors” and guided order trees with default parameters (IntegrateData). Principle component analysis (PCA) was performed on the 3,000 variable genes calculated earlier (function RunPCA). A UMAP dimensional reduction was performed on the scaled matrix using the first 25 PCA components to obtain a two-dimensional representation of cell states. Then, these defined 25 dimensionalities were used to refine the edge weights between any two cells based on Jaccard similarity (FindNeighbors), and were used to cluster cells through FindClusters functions, which implemented shared nearest neighbor modularity optimization with a resolution of 0.3, leading to 21 clusters.

To characterize clusters, the FindAllMarkers function with logfold threshold = 0.25 and minimum 0.25-fold difference and MAST test were used to identify signatures alone with each cluster. The macrophage/monocytes (clusters 0, 1, 2, 4, 6, 13, and 17)(Fig. S1 E) were selected and the top 3,000 variable features were recalculated to recluster to a higher resolution of 1. Macrophages were selected based on clusters with high expressions of known macrophage marker genes, including *Csf1r*, *C1qa*, *C1qb*, *and H2-Aa,* and confirmed by the absence of *Cd3e*, *Ms4a1*, *Krt19*, *Zbtb46*, and *Flt3*, and further confirmed by identifying DEGs associated with potential macrophage clusters, when compared to known macrophage specific marker genes. In Fig. 1 J, we reran SCTransform without regressing-out cell cycle scores to visualize proliferating macrophage clusters. In Fig. 4, L and M, monocyte clusters were removed based on expressions of monocyte markers, *Ly6c2*, *Plac8*, and *Vcan*. Macrophages were then stratified based on p21 expression into p21^High^ (top 10%) and p21^Low^ (bottom 10%), resulting in 219 of p21^High^ vs. 182 of p21^Low^ TAMs in KPC tumor, and 475 of p21^High^ vs. 526 of p21^Low^ TAMs in KP-2 orthotopic tumors. For GSEA comparisons, the log_2_ (fold-change) of all genes detected with min.pct > 0.1 and past MAST test was used as a ranking metric. GSEA was performed using GO terms, KEGG pathways, Reactome, and MSigDB gene sets with Benjamini-Hochberg FDR < 0.05 in ClusterProfiler (Wu et al., 2021). For DEGs between the two groups in each mouse PDAC model, we filtered genes with a Bonferroni-corrected p-value < 0.05 and fold-change >1.2 or <0.8.

For the mouse dataset (Hosein et al., 2019), the filtered feature barcode matrices, containing KIC, KPC, and KPFC, were processed similarly with major cell types annotated in Fig. 3 B. Macrophages were then selected and stratified based on p21 expressions into p21^High^ (top 10%) and p21^Low^ (bottom 10%), resulting in 263 of p21^High^ TAMs vs. 237 of p21^Low^ TAMs.

For p21^CE^ and p21^WT^ comparisons, the filtered feature barcode matrix was processed similarly, ending with 16,931 cells for p21^WT^ tumors, and 9,519 cells for p21^CE^ tumors. Cell cycle scores and the corresponding cell cycle phase for each cell were calculated and assigned after SCTransform based on the expression signatures for S and G2/M associated genes (CellCycleScoring). The top 3,000 variable genes, 25 dimensionalities, and resolution of 0.3 generated 19 clusters (Fig. S4, A and B), including 16,093 cells for p21^CE^ tumors and 8,996 cells for p21^WT^ tumors. Each population, including macrophages (clusters 1, 3, 5, 12, 15, and 18), monocytes (cluster 2), DCs (clusters 4, 11, 9, and 16), neutrophils (cluster 14), and eosinophils (cluster 0) were subsetted, at 15 dimensionalities and resolutions of 1 to generate Fig. 6 C and Fig. 7 A and Fig. S4 E. Cell cycle effects were also regressed-out when subsetting on each cell type, except for macrophages. DEGs with minimum percentage > 0.1, a Bonferroni-corrected p-value < 0.05. and fold-change > 1.3 or < 0.75 were considered significant. The log_2_ (fold-change) of all genes detected with minimum percentage > 0.1 and past MAST tests were used as a ranking metric for GSEA analysis. Gene sets with FDR < 0.05 were considered significant.

### Human scRNAseq data analysis

For the human dataset (Peng et al., 2019), cells with greater than 15% mitochondrial genes were retained and cells that expressed less than 500 genes were excluded. SCTransform with default parameters was used on each individual sample to normalize and scale the expression matrix against sequence depth and percentage of mitochondrial genes. Cell cycle scores and the corresponding cell cycle phase for each cell were calculated, then assigned after SCTransform based on the expression signatures for S and G2/M genes (CellCycleScoring). The differences between S phase scores and G2/M scores were regressed-out by SCTransform on individual samples. Variable features were calculated for every sample in the dataset independently and ranked based on the number of samples they were independently identified (SelectIntegrationFeatures). The top 3,000 shared variable features were used for PCA. The calculated PCA embedding of each cell was then used as an input for the soft k-means clustering algorithm. Briefly, through iteration, the algorithm designated the cluster-specific centroids and cell-specific correction factors corresponding to batch effects. The correction factors were used to assign cells into clusters until the assignment was stable (RunHarmony). Afterwards, similar steps were taken; UMAP reduction used the first 20 PCA components and FindClusters with a resolution of 0.3, leading to 12 clusters (Fig. 3 D). Immune cell clusters (3, 4, 9, and 10) were reclustered, reintegrated (RunHarmony), and UMAP reduction was used with a resolution of 0.5 to generate 11 clusters. The clusters were further grouped into NKT cells, T_regs_, T cells, Myeloid cells, and B cells in Fig. 1, F and G.

### The mpIHC

Mouse tissues were fixed in 10% formalin for 24 h and embedded in paraffin after graded ethanol dehydration. Embedded tissues were sectioned into 6-µm sections and loaded into BOND R_X_m (Leica Biosystems, Wetzlar, Germany) for a series of staining including F4/80, p21, PDPN, Ki67, and CK19. Based on antibody host species, default manufacturer protocols were used (IntenseR and Polymer Refine), containing antigen-retrieval with citrate buffer, goat serum and peroxide block, primary antibody incubation, post-primary incubation, and chromogenically visualized with an AEC substrate (Abcam, Cambridge, UK). Between each two cycles of staining, the slides were manually stained for hemoxylin and eosin, then scanned by Axio Scan.Z1 (Zeiss, Jena, Germany). The slides were then destained by a gradient of ethanol plus a 2% hydrochloride wash and blocked with extra avidin/biotin (Vector Laboratories, Burlingame, CA, USA) and a Fab fragment block (Jackson Laboratory, Bar Harbor, ME, USA).

Images of the same specimen but different stains were cropped into multiple segments by Zen software (Zeiss). Each segment was then deconvoluted (Deconvolution, v.1.0.4; Indica Labs, Albuquerque, NM, USA) for individual staining and fused using HALO software (Zeiss) with the default manufacturer’s settings. Markers of interest were pseudo-colored and quantified through the High plex FL, v.4.0.3 algorithm (Indica Labs).

## Acknowledgements

We thank the Washington University Transgenic Vector Center for generating constructs of p21^CE^ mice. We thank the Washington University Center for Cellular Imaging for imaging experiments. We thank The CHiiPs Immunomonitoring Laboratory for CyTOF experiments. We thank the Flow Cytometry & Fluorescence Activated Cell Sorting Core for sorting and flow cytometry experiments. We thank the Genome Technology Access Center for scRNAseq and microarray experiments.

## Online supplemental material

**Fig. S1** examine mpIHC staining and identifies cell types in Human CyTOF and murine scRNAseq analysis. **Fig. S2** supports siRNA knockdown of p21 in BMDMs *in vitro,* and shows p21 expression in TAMs and its connection to cell-cycle states. Representative images used to evaluate mpIHC staining of murine PDAC tissues are included. **Fig. S3** demonstrates flow cytometry and complete blood count analysis of the immune compositions in non-tumor bearing p21^CE^ and p21^WT^ mice, and in tumor bearing p21^CE^ and p21^WT^ mice. **Fig. S4** identifies major clusters in scRNAseq analysis performed on tumor bearing p21^CE^ and p21^WT^ mice. It also provides bar plots for macrophage depletion experiment. **Fig. S5** shows the GSEA results when comparing TAMs from p21^CE^ to p21^WT^. **Table S1** includes the top 50 differentially expressed genes in p21-deprived BMDMs cultured in tumor conditioned medium. **Table S2** includes all siRNA sequences used in the current paper. **Table S3** includes antibodies used for flow cytometry. **Table S4** includes all antibodies used for mpIHC. **Tables S5** and **S6** include all antibodies used for CyTOF. **Table S7** lists all primers used for qPCR. **Table S8** lists all organisms and strains used. **Table S9** lists all softwares and algorithms. Chemicals and recombinant proteins used could be found in **Table S10**. **Table S11** shows all the commercial assays used.

---

**Figure S1.**
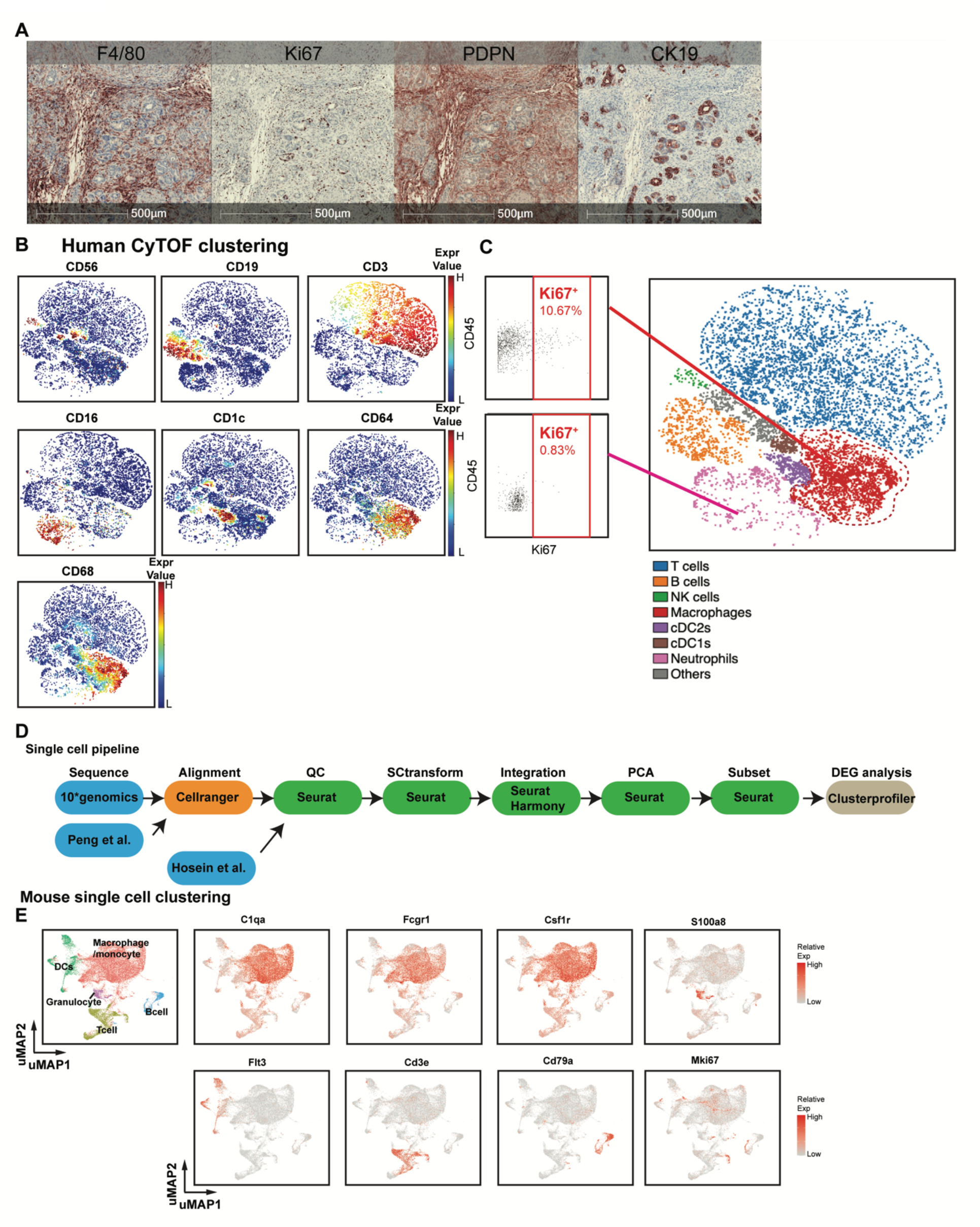
**(A)** Representative image of multiplex immunochemistry (mpIHC) staining for F4/80^+^, Ki67^+^, PDPN^+^, and CK19^+^ cells in p48^−^Cre^+^/LSL-Kras^G12D^/p53^flox/flox^ (KPC) genetically engineered mouse model (GEMM) pancreatic ductal adenocarcinoma (PDAC) tumors. Individual staining of the same samples were deconvoluted and merged through HALO software. Markers of interest were pseudo-colored and quantified through the Indica Labs-Highplex FL v.4.0.3 algorithm; n = 6. **(B)** Representative tSNE plots of human pancreatic adenocarcinoma (PDAC) samples, displaying markers used for identifying major cell types, CD56^+^ for natural killer cells, CD19^+^CD3^+^ for T cells, CD16^+^ for neutrophils, CD68^+^CD64^+^CD14^+^ for macrophages, CD1c^+^ for cDC2, and CD141^+^ for cDC1 cells; n = 9 PDAC patients. **(C)** Representative Ki67^+^ gating in macrophage and neutrophil clusters. **(D)** Schematic of the scRNAseq analysis pipeline. Details of each step for the specific dataset are listed in **Methods.** **(A) (E)** UMAP plots of integrated sorted murine CD45^+^ cells (from normal pancreas, pancreatic tissues from KPC GEMMs and orthotopic PDAC tumors) with normalized expression levels of key genes across subpopulations.

**Figure S2.**
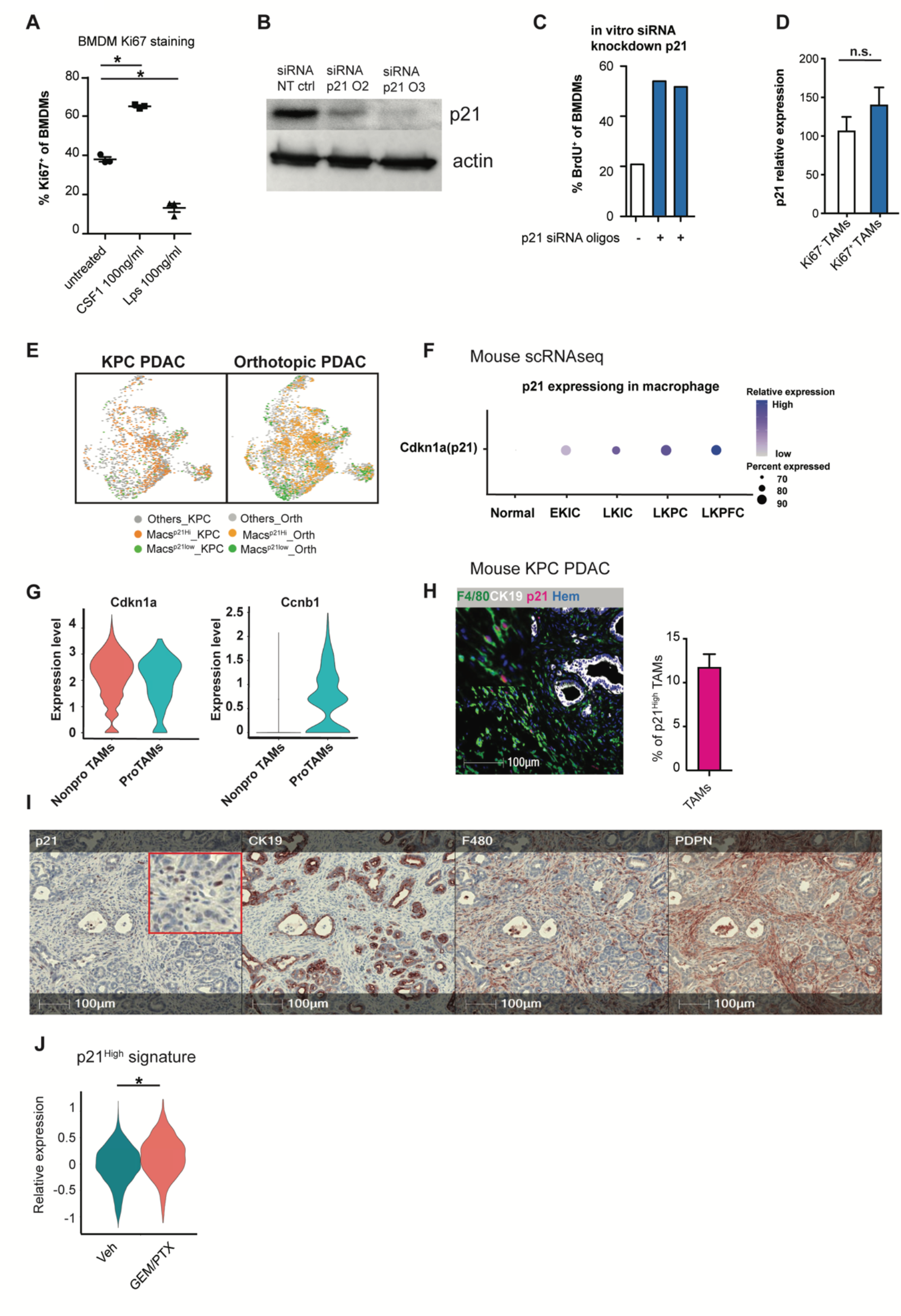
**(A)** Dot plot displaying the percentage of Ki67^+^ macrophages in bone marrow-derived macrophages (BMDMs) after colony stimulating factor-1 (CSF1) or lipopolysaccharide treatment for 24 h; n = 3/group. **(B)** Immunoblot showing expression of p21 in BMDMs after treatment with non-targeting siRNA or siRNA targeting for p21 in the presence of CSF1 for 24 h. Experiments were repeated in more than three independent repeats, and also included tumor conditioned-medium (TCM) treatment or were cultured with fibroblasts in Transwell assays. **(C)** Bar plot displaying quantification of BrdU^+^ BMDMs in **B.** The 5-bromo-2’-deoxyuridine (BrdU) was pulsed for 20 h. The experiments were repeated three times with three different siRNA oligonucleotides. **(D)** Bar plot showing the expression levels of p21 in Ki67^+^ and Ki67^−^ tumor-associated macrophages (TAMs) identified in **Fig. S1 C**; n = 9. **(E)** UMAP displaying p21^High^ and p21^Low^ macrophages in p48^−^Cre^+^/LSL-Kras^G12D^/p53^flox/flox^ (KPC) pancreatic ductal adenocarcinoma (PDAC) tumors and orthotopic KP-2 tumors. **(F)** Dot plot showing Cdkn1a (p21) gene expressions in the normal pancreas and pancreatic tissue from EKIC, LKIC, LKPC, and LKPFC genetically engineered mouse models (Hosein et al., 2019). **(G)** Violin plot of the expressions of p21 and Ccnb1 in non-proliferating and proliferating macrophages in the mouse scRNAseq dataset from the KPC, orthotopic KP-2, and normal pancreas in **Fig. 1 L**. **(H)** Representative image of multiplex immunochemistry (mpIHC) for F4/80^+^ macrophages, CK19^+^ tumor cells, and p21^+^ cells with quantification of p21^+^ macrophages from KPC PDACs; n = 8. **(I)** Representative mpIHC images of KPC mouse PDACs displaying p21, CK19, F4/80, and Pdpn staining; n = 8. **(J)** Violin plot displaying the expressions of p21^High^ signature scores, identified in **Fig. 4 M**, in tumor-associated macrophages from KPC mice 24 h after gemcitabine and paclitaxel (GEM/PTX) or dimethyl sulfoxide treatment. All graphs are expressed as the mean ± SEM. n.s., not significant; *p < 0.05. All *in vitro* assays were consistent across more than two independent repeats. For comparisons between any two groups, Student’s two-tailed *t*-test was used, except for **J** where the Bonferroni-corrected adjusted p-value was used.

**Figure S3.**
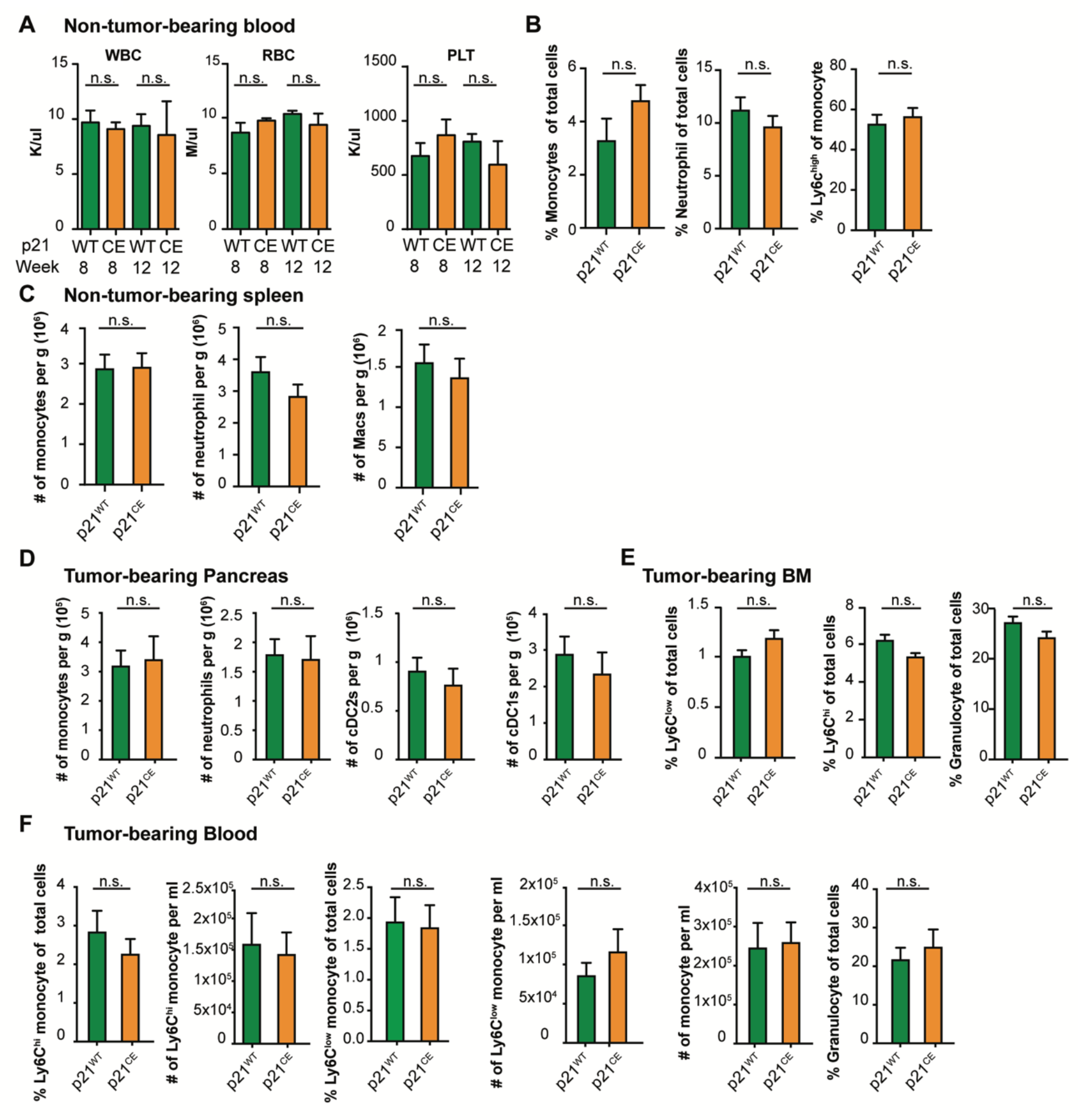
**(A)** Quantification of white blood cells, red blood cells, and platelets in non-tumor-bearing p21^WT^ and p21^CE^ mice at weeks 8 and 12; n = 3–4 mice/group. **(B)** Flow cytometry quantification of total monocytes, neutrophils, and Ly6C^hi^ monocytes in blood of non-tumor-bearing p21^WT^ and p21^CE^ mice; n = 7–9 mice/group. **(C)** Flow cytometry quantification of monocytes, neutrophils, and macrophages in the spleens of 8–12 weeks p21^WT^ and p21^CE^ non-tumor-bearing mice; n = 7–9 mice/group. **(D)** Flow cytometry analysis of the number of monocytes, neutrophils, cDC2s, and cDC1s in the pancreas of p21^CE^ and p21^WT^ mice bearing orthotopic KP-2 tumors; n = 6 mice/group. **(E)** Flow cytometry quantification of Ly6C^hi^ monocytes, Ly6C^low^ monocytes, and granulocytes in the bone marrow of tumor-bearing p21^CE^ and p21^WT^ mice; n = 6 mice/group. **(F)** Flow cytometry quantification of myeloid cells in the blood of tumor-bearing p21^CE^ and p21^WT^ mice; n = 6 mice/group. All graphs are expressed as the mean ± SEM. n.s., not significant; *p < 0.05. For comparisons between any two groups, the Student’s two-tailed *t*-test was used.

**Figure S4.**
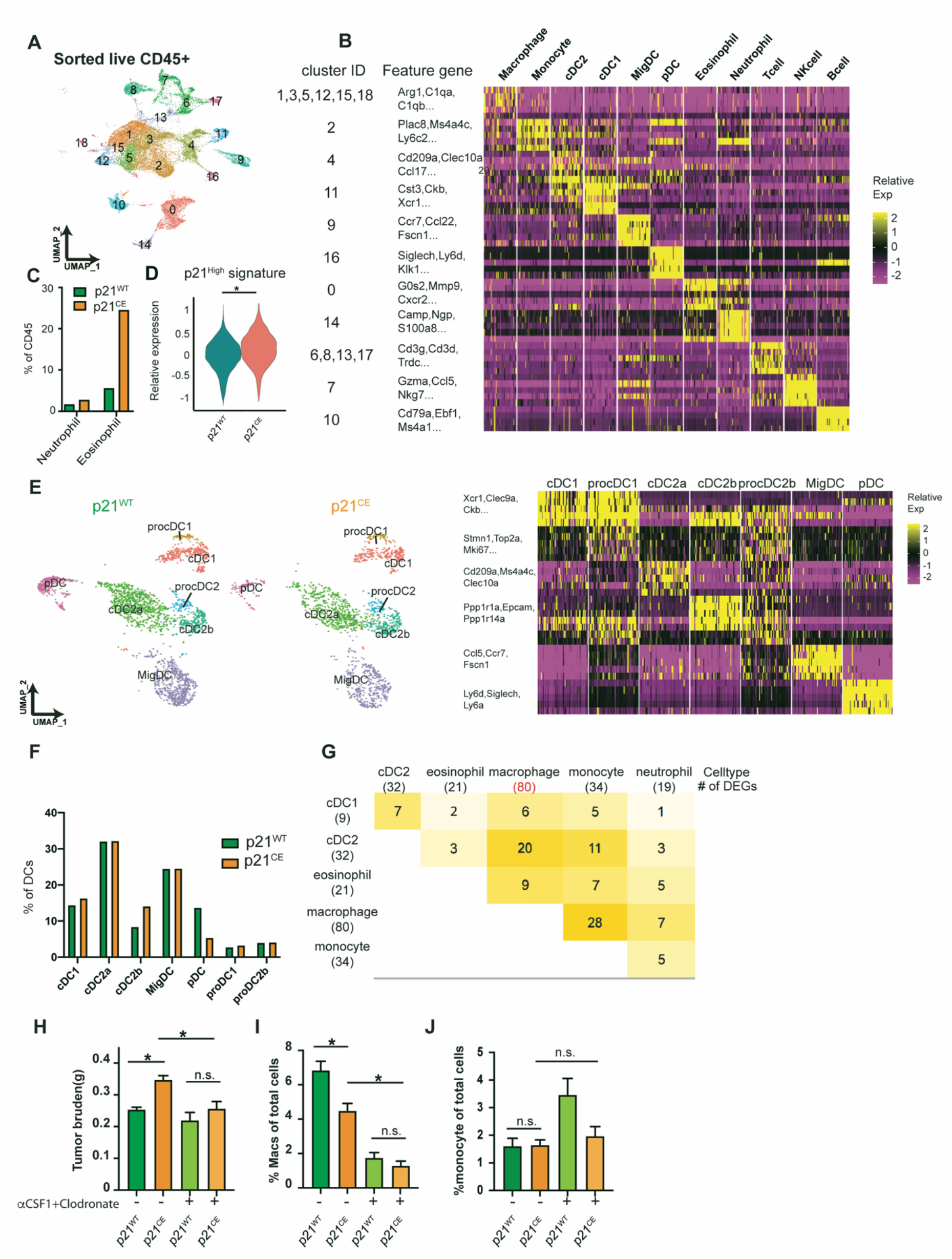
**(A)** UMAP plot of all sorted CD45^+^ cell clusters on merged objects from p21^CE^ and p21^WT^ KP-2 orthotopic tumor-bearing mice. Three mice were pooled for each genotype. **(B)** Heat map listing all clusters in **A** and corresponding cell type annotations and key gene expressions. **(C)** Bar plot displaying the percentages of neutrophils and eosinophils in p21^WT^ and p21^CE^ tumor-bearing mice. **(D)** Violin plot displaying the expression levels of p21^High^ signature scores, identified in **Fig. 4 M**, in TAMs from p21^CE^ and p21^WT^ mice. *Wilcox adjusted p.value < 0.05. **(E)** UMAP plot of the reclustered DC populations in **Fig. 6 A**, annotated with cell type and associated key gene expressions in the heat map (right). **(F)** Quantification of major DC populations identified in **D** from p21^CE^ tumors when compared with p21^WT^ tumors. **(G)** Heat map showing the number of shared differentially-expressed genes (DEGs) between two genotypes in each cell population, including macrophage and close lineages. The number of DEGs for each single cell population when comparing p21^CE^ to p21^WT^ was listed in the parenthesis below. **(H-J)** Bar plot showing the tumor burden, percentages of tumor-associated macrophages and monocytes in p21^WT^ and p21^CE^ mice bearing orthotopic KP-2 tumors with or without colony stimulating factor-1 and clodronate treatment; n = 8–10 mice/group. n.s., not significant; *p < 0.05. For comparisons between any two groups, the Student’s two-tailed *t*-test was used.

**Figure S5.**
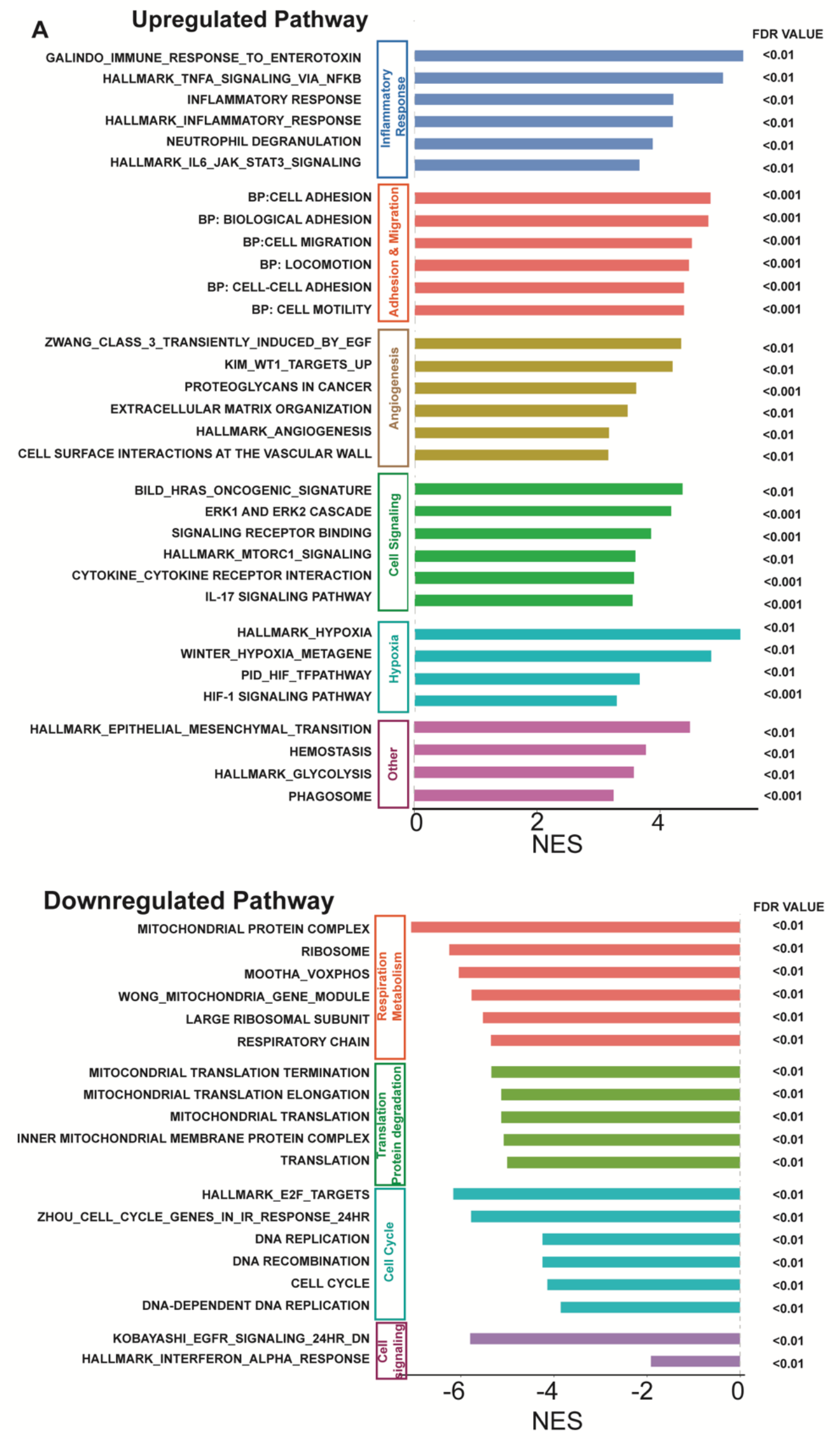
**(A)** Bar plot showing significantly upregulated and downregulated pathways identified by GSEA in tumor-associated macrophages from p21^CE^ compared with p21^WT^ mice. The pathways were grouped into biological functions with a false discovery rate < 0.01.

**Table S1:** Array top regulated genes sip21 vs. siNT; n = 3 each.

| GeneSymbol | Fold Change<br>(sip21o2 vs siNT) | adjusted p | Fold Change<br>(sip21o3 vs siNT) | adjusted p |
| --- | --- | --- | --- | --- |
| Hfm1 | -9.000967 | 4.17E-06 | -9.551253 | 3.30E-06 |
| Olfr356 | -6.310875 | 0.000277 | -6.7342 | 0.000213 |
| Cnn1 | -4.243251 | 2.60E-05 | -4.088136 | 3.23E-05 |
| Supt3 | -3.3844 | 6.81E-05 | -5.437286 | 4.28E-06 |
| Cdkn1a | -2.804869 | 1.44E-07 | -5.631165 | 1.15E-09 |
| Ear7 | -2.404938 | 1.27E-06 | -2.886804 | 2.29E-07 |
| Cdkn1a | -2.396397 | 4.53E-07 | -3.167768 | 3.54E-08 |
| Spint1 | -1.945373 | 0.000275 | -2.055021 | 0.00015 |
| Spint1 | -1.826354 | 5.97E-05 | -1.544564 | 0.000716 |
| Slc36a2 | -1.781509 | 2.15E-05 | -1.581507 | 0.00014 |
| Ldhb | -1.71729 | 0.000103 | -1.817892 | 4.60E-05 |
| 0610009E02Rik | -1.704009 | 0.000111 | -1.568371 | 0.000405 |
| Hpgd | -1.689703 | 7.12E-05 | -1.759566 | 3.89E-05 |
| Rcbtb2 | -1.67373 | 4.01E-05 | -1.513442 | 0.000225 |
| Gm9733 | -1.579973 | 0.000145 | -1.708255 | 4.09E-05 |
| Aldoc | -1.547172 | 6.71E-06 | -1.556962 | 5.93E-06 |
| Ppp1r9a | -1.54189 | 5.78E-05 | -1.602184 | 2.88E-05 |
| Sult1a1 | -1.538296 | 0.000123 | -1.649467 | 3.66E-05 |
| Cib2 | -1.522033 | 0.000125 | -1.524215 | 0.000121 |

Top 50 upregulated genes
| GeneSymbol | Fold-change<br>(sip21o2 vs TCM) | adjusted p | Fold-change<br>(sip21o3 vs TCM) | adjusted p |
| --- | --- | --- | --- | --- |
| Rrm2 | -6.30647 | 3.17E-11 | -7.95356 | 1.02E-11 |
| Cdkn3 | -6.28505 | 8.43E-10 | -6.7755 | 5.76E-10 |
| Pif1 | -6.25111 | 2.88E-07 | -8.06132 | 8.74E-08 |
| Cxcl9 | -5.11269 | 1.94E-05 | -5.20062 | 1.78E-05 |
| Hmmr | -4.85418 | 3.20E-05 | -5.03176 | 2.65E-05 |
| Hmmr | -4.31765 | 9.85E-05 | -4.58722 | 7.12E-05 |
| Mastl | -4.07957 | 9.67E-07 | -4.21229 | 7.90E-07 |
| Casc5 | -3.96834 | 4.20E-05 | -5.70901 | 5.80E-06 |
| Pif1 | -3.89545 | 2.18E-05 | -4.68433 | 7.38E-06 |
| Fancd2 | -3.83321 | 2.11E-05 | -4.92167 | 4.91E-06 |
| Esco2 | -3.67204 | 6.75E-06 | -5.21945 | 8.15E-07 |
| D17H6S56E-5 | -3.60167 | 1.85E-08 | -4.03994 | 8.27E-09 |
| Mastl | -3.59825 | 4.18E-06 | -4.83099 | 6.63E-07 |
| Kif2c | -3.54186 | 1.80E-06 | -4.37275 | 4.51E-07 |
| Nek2 | -3.52283 | 2.45E-08 | -4.65013 | 3.78E-09 |
| 2010110K18Rik | -3.49874 | 1.67E-05 | -4.71255 | 2.65E-06 |
| Cdca2 | -3.46713 | 0.000389 | -4.92359 | 5.65E-05 |
| Dlgap5 | -3.41372 | 9.11E-08 | -4.60248 | 1.20E-08 |
| Xkr5 | -3.39351 | 0.000359 | -4.19719 | 0.000105 |
| Foxm1 | -3.38906 | 8.91E-08 | -4.27792 | 1.76E-08 |
| Prc1 | -3.34136 | 3.54E-09 | -4.5658 | 4.01E-10 |
| Fbxo48 | -3.29161 | 7.43E-06 | -4.39264 | 1.10E-06 |
| Prc1 | -3.25269 | 3.85E-07 | -4.169 | 6.68E-08 |
| Sgol1 | -3.21025 | 8.10E-07 | -3.98796 | 1.72E-07 |
| BC030867 | -3.20821 | 2.85E-08 | -4.46017 | 2.75E-09 |
| Depdc1a | -3.19559 | 0.000114 | -3.78934 | 3.79E-05 |
| Ckap2l | -3.17264 | 0.000106 | -4.27054 | 1.62E-05 |
| Kifc5b | -3.13167 | 0.000538 | -4.41533 | 7.18E-05 |
| Cenpf | -3.11857 | 7.64E-05 | -3.32338 | 4.92E-05 |
| Nusap1 | -3.0947 | 2.82E-09 | -4.11654 | 3.34E-10 |
| Kif4 | -3.09329 | 0.000319 | -3.63404 | 0.000115 |
| Kifc1 | -3.0841 | 5.61E-05 | -3.56142 | 2.08E-05 |
| Kif18b | -3.08083 | 1.24E-05 | -4.31059 | 1.26E-06 |
| Prc1 | -3.07972 | 0.0001 | -3.4257 | 4.83E-05 |
| Rad51 | -3.07141 | 6.81E-08 | -3.88828 | 1.15E-08 |
| Pbk | -3.06893 | 7.85E-06 | -3.44022 | 3.38E-06 |
| Aspm | -3.06868 | 1.39E-06 | -4.04615 | 1.89E-07 |
| Sgol1 | -3.0153 | 1.27E-07 | -4.3219 | 9.18E-09 |
| Rad51ap1 | -3.01418 | 2.51E-08 | -4.12435 | 2.40E-09 |
| Ccna2 | -3.01189 | 1.32E-06 | -3.30825 | 6.34E-07 |
| Rad51ap1 | -2.99682 | 1.99E-06 | -3.89215 | 2.92E-07 |
| Efcab6 | -2.99481 | 2.19E-05 | -2.63158 | 6.16E-05 |
| Fam64a | -2.98438 | 8.13E-08 | -4.08292 | 7.76E-09 |
| Shcbp1 | -2.9764 | 0.000173 | -3.27637 | 8.95E-05 |
| Ccnb2 | -2.97509 | 1.24E-08 | -3.76131 | 1.98E-09 |
| Ccnb1 | -2.97083 | 2.59E-09 | -4.04031 | 2.45E-10 |
| Cdc20 | -2.96576 | 7.65E-08 | -3.80712 | 1.12E-08 |
| Ccnb1 | -2.96254 | 2.22E-07 | -3.57911 | 5.06E-08 |
| Anln | -2.95577 | 1.58E-05 | -3.15593 | 9.56E-06 |

**Table S2:**
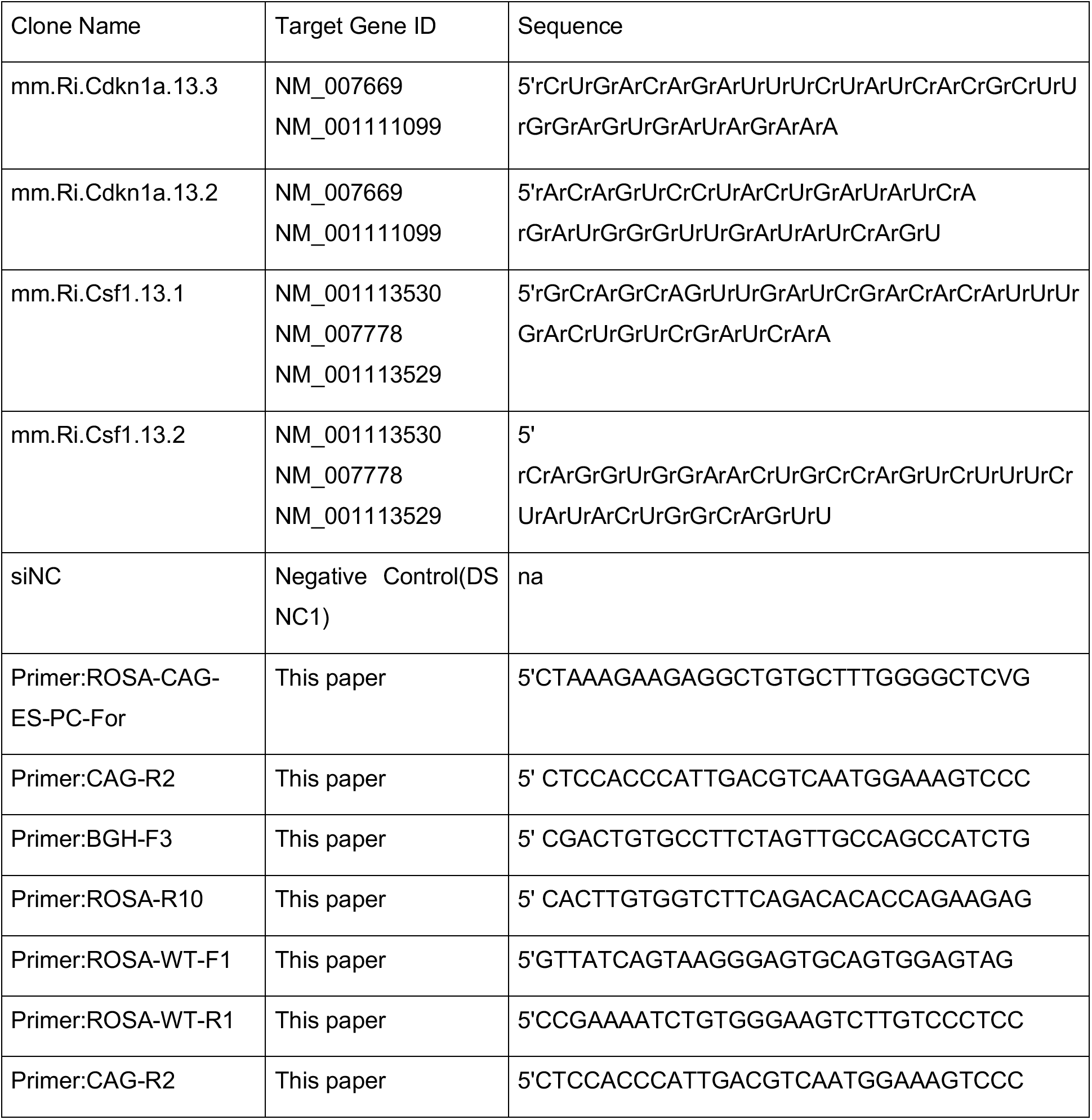
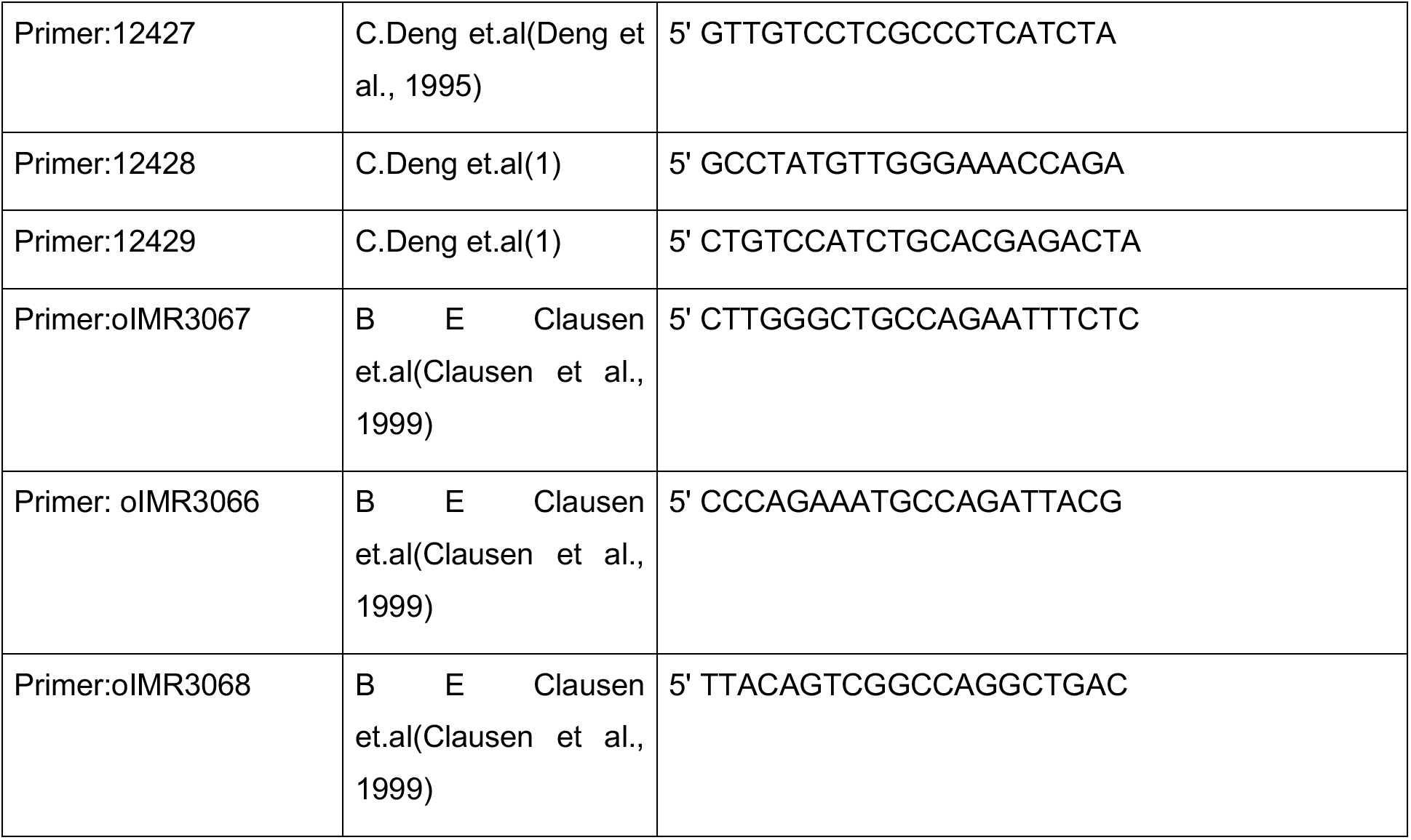
Sequences of siRNAs targeting p21 and CSF1

**Table S3:** Antibody list for flow cytometry

| Name | Identifier | Clone# | Company | Dilution |
| --- | --- | --- | --- | --- |
| CD45 | RRID:AB_469625 | 30-F11 | eBioscience | 1:400 |
| CD3e | RRID:AB_469315 | 145-2C11 | eBioscience | 1:200 |
| CD4 | RRID:AB_464900 | RM4-4 | eBioscience | 1:200 |
| CD8a | RRID:AB_2732919 | 53-6.7 | BD Biosciences | 1:200 |
| Foxp3 | RRID:AB_11218094 | FJK-16s | eBioscience | 1:100 |
| CD19 | RRID:AB_1659676 | eBio1D3 | eBioscience | 1:200 |
| CD11b | RRID:AB_657585 | M1/70 | eBioscience | 1:400 |
| CD11c | RRID:AB_1548652 | N418 | eBioscience | 1:200 |
| Ly6C | RRID:AB_1518762 | HK1.4 | eBioscience | 1:400 |
| Ly6G | RRID:AB_1186104 | 1A8 | BioLegend | 1:400 |
| F4/80 | RRID:AB_468798 | BM8 | eBioscience | 1:400 |
| MHCII | RRID:AB_1272204 | M5/115.15.2 | eBioscience | 1:400 |
| CD24 | RRID:AB_464985 | 30-F1 | eBioscience | 1:200 |
| CD44 | RRID:AB_1272246 | IM7 | eBioscience | 1:200 |
| CD62L | RRID:AB_11125577 | MEL-14 | BioLegend | 1:100 |
| Flt3 | RRID:1B_1877218 | A2F10 | BioLegend | 1:20 |
| CD115 | RRID:AB_467428 | AFS98 | eBioscience | 1:50 |
| B220 | RRID:AB_396673 | RA3-6B2 | eBiosciences | 1:100 |
| gdTCR | RRID:AB_842756 | eBioGl3 | eBioscience | 1:200 |
| Nk1.1 | RRID:AB_467736 | PK136 | eBioscience | 1:100 |
| Sca1 | RRID:AB_467778 | D7 | BioLegend | 1:100 |
| Cd49b | RRID:AB_395093 | DX5 | eBioscience | 1:200 |

**Table S4:** Antibody list for multiplex immunochemistry (mpIHC) and immunoblotting

| Name | Identifier | Clone# | Company | Dilution |
| --- | --- | --- | --- | --- |
| P21 | Ab188224 | EPR18021 | Abcam | 1:200 |
| Ck19 | RRID:AB_469315 | 145-2C11 | eBioscience | 1:200 |
| Ki-67 | 12202 | D3B5 | Cell Signaling | 1:400 |
| Podoplanin | Ab11936 | RTD4E10 | Abcam | 1:800 |
| F4/80 | 70076 | D2S9R | Cell Signaling | 1:200 |
| pp65 | 3033 | 93H1 | Cell Signaling | 1:500 |
| Cyclin d1 | 2926 | DCS6 | Cell Signaling | 1:500 |
| c-Myc | 13987 | D3N8F | Cell Signaling | 1:500 |
| P27 | 3686 | D69C12 | Cell Signaling | 1:500 |

**Table S5:** Antibody list for human mass cytometry time of flight

| REAGENT or RESOURCE | SOURCE | IDENTIFIER |
| --- | --- | --- |
| anti-human CD11b (ICRF44) | Fluidigm | #3209003B |
| anti-human CD11c (Bu15) | Fluidigm | #3159001B |
| anti-human CD14 (M5E2) | Fluidigm | #3160001B |
| anti-human CD141 (1A4) | Fluidigm | #3173002B |
| anti-human CD15 (W6D3) | Fluidigm | #3164001B |
| anti-human CD16 (3G8) | Fluidigm | #3148004B |
| anti-human CD163 (GHI/61) | Fluidigm | #3154007B |
| anti-human CD19 (HIB19) | Fluidigm | #3142001B |
| anti-human CD192 (CCR2) (K036C2) | Fluidigm | #3153023B |
| anti-human CD1c (L161) | BioLegend | #331502 |
| anti-human CD20 (2H7) | Fluidigm | #3147001B |
| anti-human CD206 (MMR) (15-2) | Fluidigm | #3168008B |
| anti-human CD24 (ML5) | Fluidigm | #3166007B |
| anti-human CD3 (UCHT1) | BioLegend | #300402 |
| anti-human CD32 (FUN-2) | Fluidigm | #3169020B |
| anti-human CD34 (581) | Fluidigm | #3149013B |
| anti-human CD38 (HIT2) | Fluidigm | #3167001B |
| anti-human CD40 (5C3) | Fluidigm | #3165005B |
| anti-human CD45 (HI30) | Fluidigm | #3089003B |
| anti-human CD54 (HA58) | Fluidigm | #3170014B |
| anti-human CD56 (NCAM16.2) | Fluidigm | #3176008B |
| anti-human CD64 (10.1) | Fluidigm | #3146006B |
| anti-human CD68 (Y1/82A) | Fluidigm | #3171011B |
| anti-human CD80 (2D10.4) | Fluidigm | #3162010B |
| anti-human CD81 (5A6) | Fluidigm | #3145007B |
| anti-human CD82 (ASL-24) | Fluidigm | #3158025B |
| anti-human CD86 (IT2.2) | Fluidigm | #3150020B |
| anti-human CX3CR1 (2A9-1) | Fluidigm | #3172017B |
| anti-human CXCR4 (12G5) | Fluidigm | #3175001B |
| anti-human HLA-DR (L243) | Fluidigm | #3174001B |
| anti-human Ki-67 (B56) | Fluidigm | #3161007B |
| Anti-human PCNA(PC10) | Abcam | Ab29 |
| Anti-human p21(12D1) | CellSignal | 2947 |

**Table S6:** Antibody list for mouse mass cytometry time of flight (CyTOF)

| REAGENT or RESOURCE | SOURCE | Catalog |
| --- | --- | --- |
| anti-mouse CD44(IM7) | Leinco | C382 |
| anti-mouse GITR(DTA-1) | BioXcell | BE0063 |
| anti-mouse CD25(PC61) | Leinco | C1194 |
| anti-mouse CD38(90) | eBioscience | 14-0381-82 |
| anti-mouse CD90(G7) | Biolegend | 105202 |
| anti-mouse Lag-3(C9B7W) | Leinco | L306 |
| anti-mouse CD27(LG.7F9) | eBioscience | 50-124-94 |
| anti-mouse KLRG1(2F1/KLRG1) | BioXCell | BE0201 |
| anti-mouse CD103(2E7) | Biolegend | 121402 |
| Anti-mouse CD4(GK1.5) | BioXcell | BE0003-1 |
| anti-mouse CD45(30-F11) | Fluidigm | 3089005B |
| anti-mouse CD62L(MEL-14) | Leinco | C2118 |
| anti-mouse ICOS(C398.4A) | eBioscience | 14-9949-82 |
| anti-mouse OX-40(OX-86) | BioXcell | BE0031 |
| anti-mouse PD-1(RMP1-30) | eBioscience | 14-9981-82 |
| anti-mouse TIGIT(1G9) | BioXcell | BE0274 |
| anti-mouse CD69(H1.2F3) | eBioscience | 14-0691-82 |
| anti-mouse TCRb(H57-597) | BioXcell | BE0102 |
| anti-mouse CD127(A7R34) | BioXcell | BE0065 |
| anti-mouse CD39(Duha59) | Biolegend | 143802 |
| anti-mouse NK1.1(PK136) | BioXcell | BE0036 |
| anti-mouse CD8a(53-6.7) | Leinco | C375 |
| anti-mouse TCRgd(GL3) | eBioscience | 14-5711-82 |
| anti-mouse Tim3(RMT3-23) | BioXcell | BE0115 |
| anti-mouse 4-1BB(17B5) | BioLegend | 106107 |
| anti-mouse FoxP3(FJK-16s) | eBioscience | 14-5773-82 |
| anti-mouse GATA3(TWAJ) | eBioscience | 14-9966-82 |
| anti-mouse GranzymeB(GB11) | eBioscience | MA1-80734 |
| anti-mouse CTLA-4(UC10-4B9) | eBioscience | 50-129-16 |
| anti-mouse Ki67(8D5) | Novus | NBP2-22112 |
| anti-mouse TCF1(812145) | R&D | MAB8224 |
| anti-mouse ROR-γ $\tau$(AFKJS-9) | eBioscience | 14-6988-82 |
| anti-mouse Eomes(Dan11mag) | eBioscience | 50-245-556 |
| Anti-mouse T-bet(4B10) | Biolegend | 644802 |

**Table S7:** List of qPCR primers

| Gene | Source | Assay ID |
| --- | --- | --- |
| GAPDH | Taqman | Mm99999915_g1 |
| TBP | Taqman | Mm01277042_m1 |
| HPRT | Taqman | Mm03024075_m1 |
| CCND1 | Taqman | Mm00432359_m1 |
| CCNE1 | Taqman | Mm01266311_m1 |
| CCNA2 | Taqman | Mm00438063_m1 |
| IFIT3 | Taqman | Mm01704846_s1 |
| CD40 | Taqman | Mm00441891_m1 |
| IFNA1 | Taqman | Mm03030145_gH |
| IFNB1 | Taqman | Mm00439546_s1 |
| IRF1 | Taqman | Mm01288580_m1 |
| CDKN1A | Taqman | Mm04205640_g1 |
| CSF1 | Taqman | Mm00432686_m1 |
| CXCL1 | Taqman | Mm04207460_m1 |
| CXCL2 | Taqman | Mm00436450_m1 |
| IL-1α | Taqman | Mm00439620_m1 |
| IL-1β | Taqman | Mm00434228_m1 |
| IL-6 | Taqman | Mm00446190_m1 |
| TNF-α | Taqman | Mm00443258_m1 |
| IRF4 | Taqman | Mm00516431_m1 |
| TGF-β2 | Taqman | Mm00436955_m1 |
| TGF-β3 | Taqman | Mm00436960_m1 |
| YM1 | Taqman | Mm00657889_mH |
| BATF | Taqman | Mm00479410_m1 |
| c-Myc | Taqman | Mm00487804_m1 |
| CCL2 | Taqman | Mm00441242_m1 |

**Table S8:**
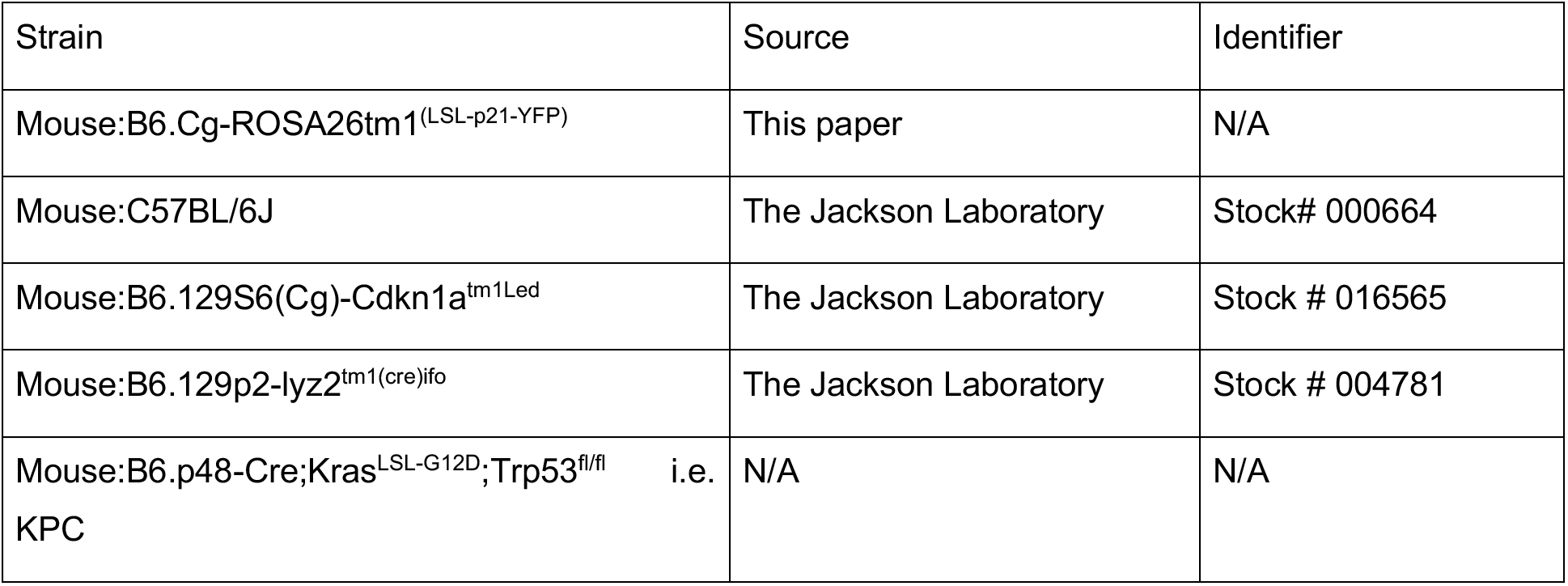
Experimental Models: Organisms/strains

**Table S9:**
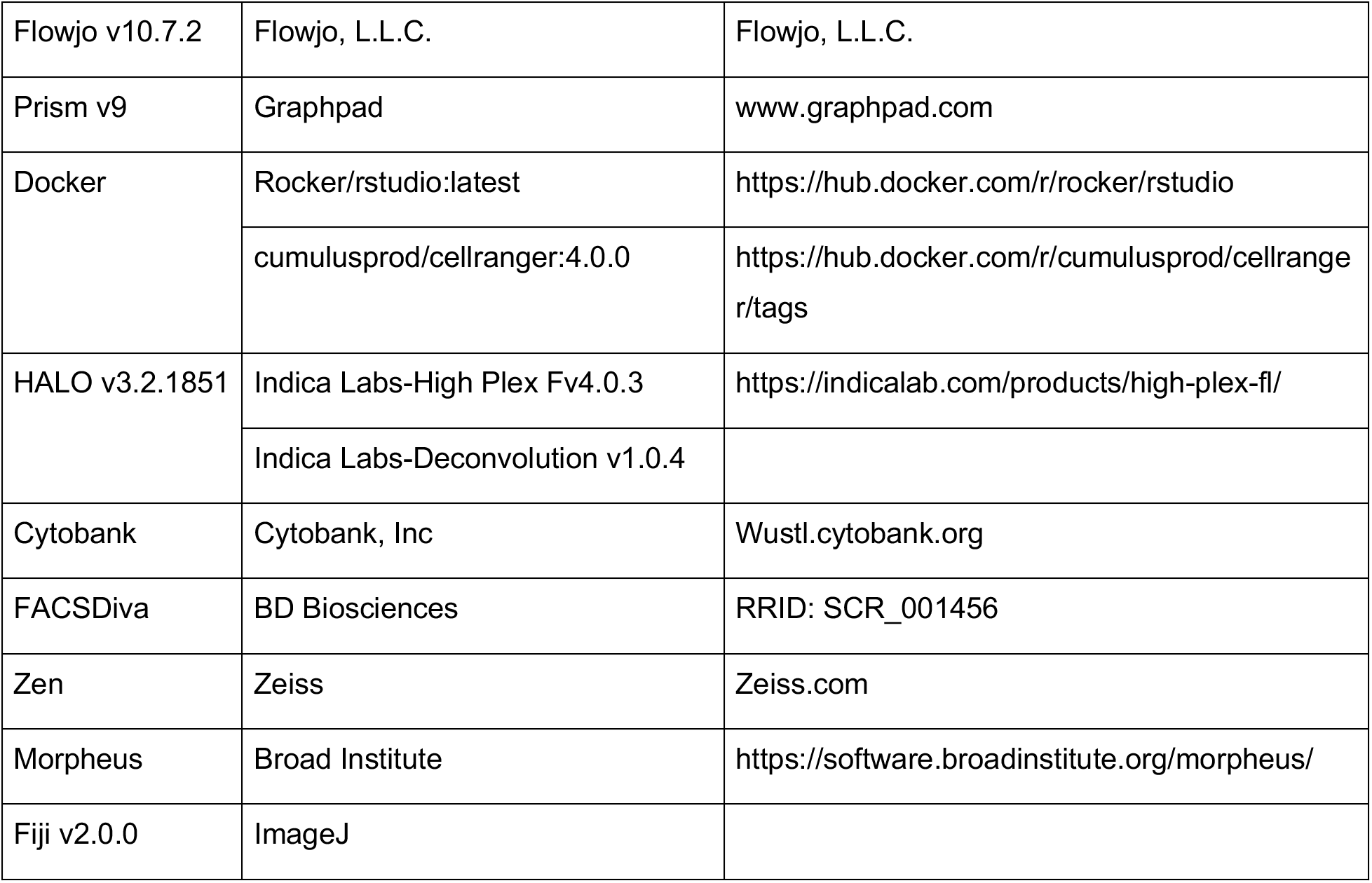
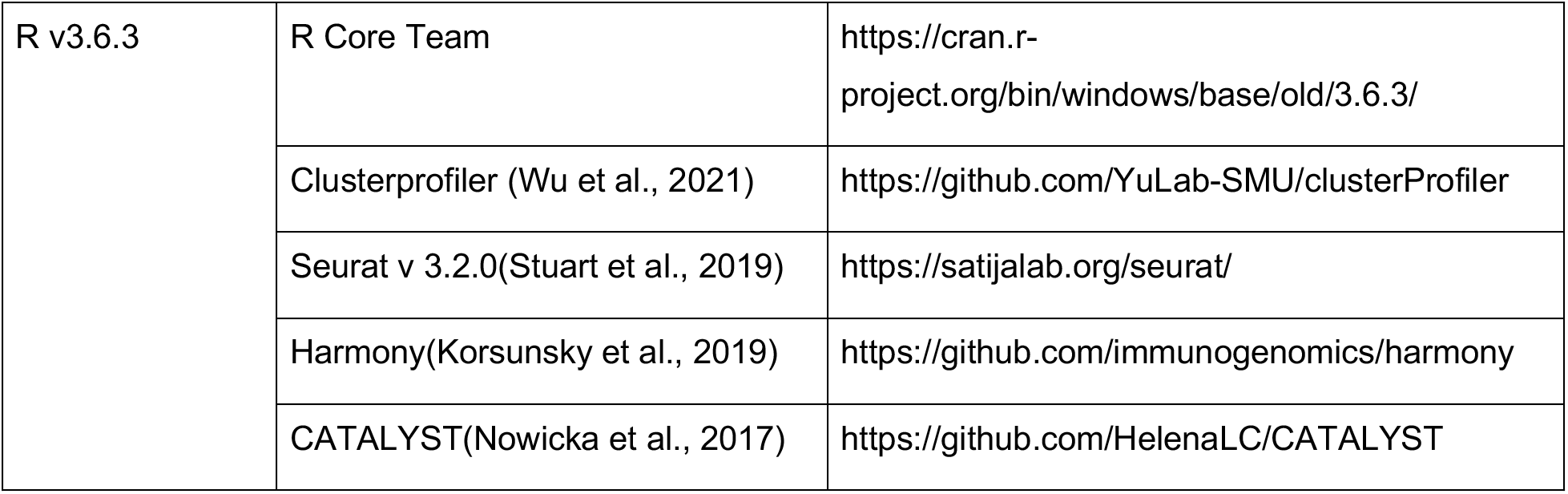
Software and Algorithms

**Table S10:**
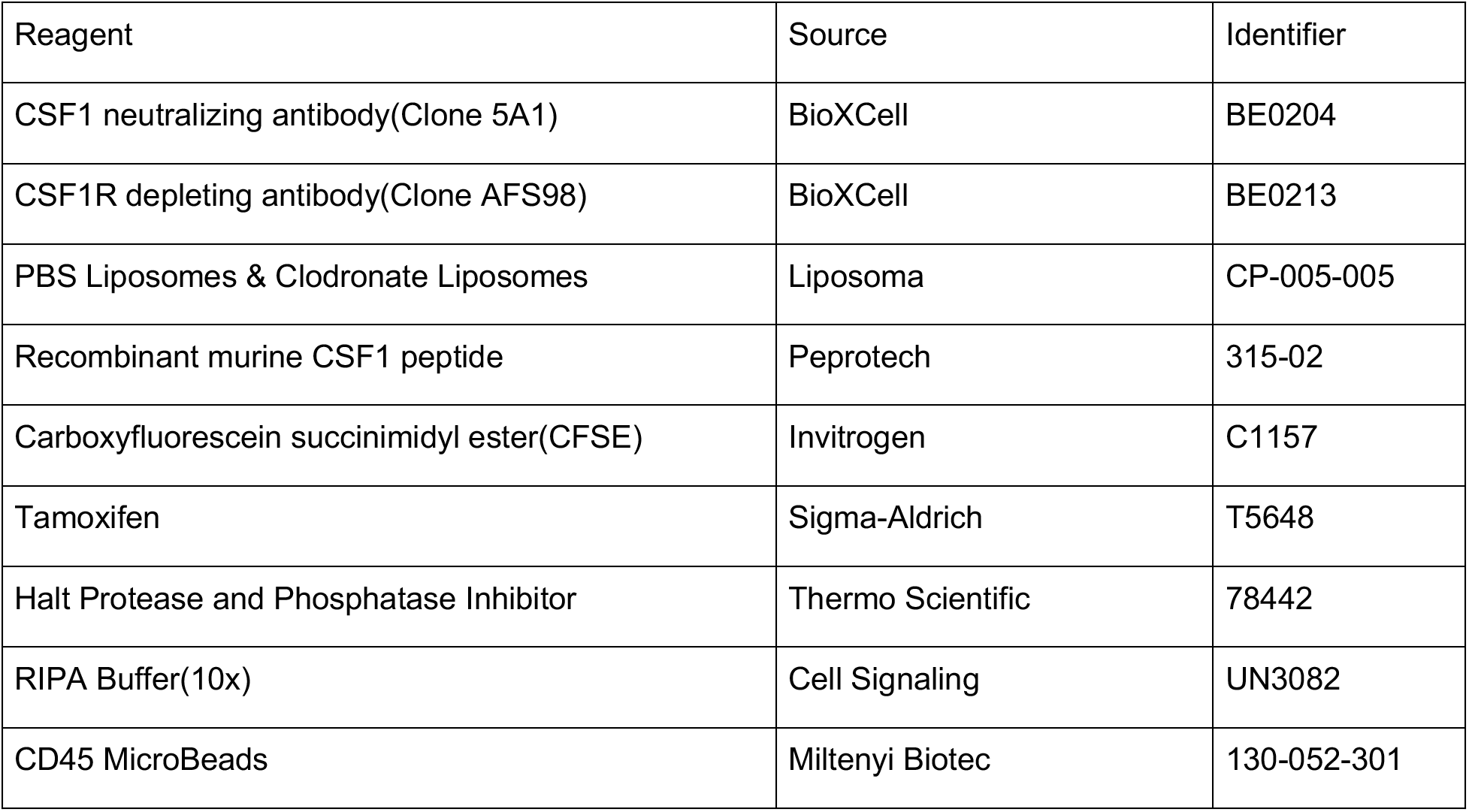
Chemicals and recombinant proteins

**Table S11:** Critical commercial assays

| Resources | Source | Catalog number |
| --- | --- | --- |
| Supersignal West Dura | Thermo | 34075 |
| FITC BrdU Flow Kit | BD Bioscience | 557891 |
| APC BrdU Flow Kit | BD Bioscience | 557892 |
| Proteome Profiler Mouse XL Cytokine Array | R&D systems | ARY028 |
| E.Z.N.A. Total RNA Kit I | Omega | R6834-02 |
| Taqman Gene Expression Master Mix | Thermo Fisher | 4370074 |
| qScript cDNA Supermix kit | Quantabio | 95048-500 |
| BOND Polymer Refine Detection Kit | Leica | DS9800 |
| BOND Polymer Refine Red Detection Kit | Leica | DS9390 |
| BOND Intense R Detection Kit | Leica | DS9263 |
| Cytofix Kit | BD Bioscience | 554655 |
| MACS LS kit | Miltenyi | 130042401 |
| Mouse Macrophage Nucleofector Kit | Lonza | VPA-1009 |
| Latex Beads-Rabbit IgG-PE Complex | Cayman | 600541 |
| Mouse M-CSF Matched Antibody Pair Kit | Abcam | Ab218788 |
| Pierce BCA Protein Assay Kit | Thermo Scientific | 23225 |
| DuoSet ELISA Mouse M-CSF | R&D system | DY416-05 |
| DuoSet Ancillary Reagent Kit2 | R&D system | DY008 |

